# Sexual antagonism in haplodiploids

**DOI:** 10.1101/2021.03.26.437233

**Authors:** Thomas J. Hitchcock, Andy Gardner, Laura Ross

## Abstract

Females and males may face different selection pressures, such that alleles conferring a benefit in one sex may be deleterious in the other. Such sexual antagonism has received a great deal of theoretical and empirical attention, almost all of which has focused on diploids. However, a sizeable minority of animals display an alternative haplodiploid mode of inheritance, encompassing both arrhenotoky, whereby males develop from unfertilized eggs, and paternal genome elimination (PGE), whereby males receive but do not transmit a paternal genome. Alongside unusual genetics, haplodiploids often exhibit social ecologies that modulate the relative value of females and males. Here we develop a series of evolutionary-genetic models of sexual antagonism for haplodiploids, incorporating details of their molecular biology and social ecology. We find that: 1) PGE promotes female-beneficial alleles more than arrhenotoky, and to an extent determined by the timing of elimination – and degree of silencing of – the paternal genome; 2) sib-mating relatively promotes female-beneficial alleles, as do other forms of inbreeding, including limited male-dispersal, oedipal-mating, and the pseudo-hermaphroditism of *Icerya purchasi*; 3) resource competition between related females relatively inhibits female-beneficial alleles; and 4) sexual antagonism foments conflicts between parents and offspring, endosymbionts and hosts, and maternal-origin and paternal-origin genes.

## Introduction

Organisms often appear remarkably well adapted to live the lives they do, as a consequence of the historical action of natural selection. Some of the best tests of our understanding of adaptation occur when organisms must make trade-offs between conflicting design objectives. Sexual antagonism is one such example, whereby genetic variants may prove beneficial to one sex but detrimental to the other. This has motivated a large body of theoretical work considering when such sexually antagonistic alleles will be able to invade (Owen 1953), how this may vary across the genome (Parsons 1961; Kidwell et al. 1977; Rice 1984; Frank and Hurst 1996; Frank and Patten 2020; Hitchcock and Gardner 2020), and how we may be able to detect such alleles from population genetic data (Cheng and Kirkpatrick 2016; Kasimatis et al. 2019; Ruzicka and Connallon 2020; Ruzicka et al. 2020). This theory has been complemented more recently by molecular and quantitative genetic studies of laboratory and wild populations, both estimating the extent of sexual antagonism, and identifying specific loci at which sexually antagonistic alleles reside (Poissant et al. 2010; Mank 2017; Rowe et al. 2018; Connallon and Matthews 2019).

Almost all this research has focused on diploid, “eumendelian” (*sensu* Normark 2006) organisms. However, a sizeable minority of animals (~15%) display an alternative, haplodiploid mode of inheritance (Normark 2003, 2006; Bachtrog et al. 2014). Haplodiploidy encompasses both arrhenotoky – whereby males develop from unfertilized eggs – and paternal genome elimination (PGE) – whereby males receive but do not transmit a paternal genome – and is employed by a diverse cast of creatures in groups as distinct as mites, nematodes, rotifers, springtails, beetles, wasps and flies. In all of these organisms, males exclusively transmit maternal-origin genes, such that reproduction of females contributes twice as much to the ancestry of future generations as does that of males. Whilst similarities in transmission genetics have drawn comparisons to X-linked genes (Kraaijeveld 2009; de la Filia et al. 2015), haplodiploids are not merely whole-organismal manifestations of X chromosomes. Firstly, mechanisms of dosage compensation – that ensure an equal balance of X-linked versus autosomal gene products between females and males – are understood to play an important role in modulating sexual antagonism in relation to the X chromosome (Hitchcock and Gardner 2020), but it is unclear whether these mechanisms should apply in the same way in relation to arrhenotokous species in which males are haploid across their entire genome. Secondly, although PGE is similar to X-linkage from a transmission perspective, this form of haplodiploidy involves males being somatically diploid through some or all of their lives (Burt and Trivers 2006; Gardner and Ross 2014), with concomitant gene dosage and dominance effects that may be expected to affect the balance between female-beneficial versus male-beneficial alleles.

Moreover, haplodiploids often exhibit characteristic social ecologies, including gregarious broods, chronic inbreeding, and strongly female-biased primary sex ratios (Hamilton 1967). An archetypal example is the date stone beetle (*Coccotrypes dactyliperda*), whereby a gravid female excavates a tunnel into a date seed and lays a large and heavily female-biased brood, her offspring then mate with each other, and her mated-daughters then leave to search for dates within which to raise their own families (Hamilton 1993; Spennemann 2019). Whilst the particular niche that these species inhabit may vary substantially – from fungal-feeding to sap- or blood-sucking – they often share a similarly viscous population structure, with small, semi-isolated subpopulations, and large amounts of inbreeding (Hamilton 1967, 1978, 1993; Normark 2006). These unusual mating systems generate peculiar patterns of within-individual and between-individual relatedness, as well as differences in the scales at which the sexes compete and cooperate. Both of these factors are known to modulate the relative genetic value of males and females in the context of sex allocation (Taylor 1981; Frank 1986b; Nagelkerke and Sabelis 1996; West 2009), and thus might also be expected to alter the outcome of sexually antagonistic selection.

Here we develop a series of evolutionary-genetic models of sexual antagonism for haplodiploids, incorporating details of both their molecular biology and social ecology. We first consider how the genetic asymmetries found in haplodiploids are expected to alter the fate of sexually antagonistic alleles, and how this is modified by variation in the timing and expression of the paternal-genome. We then explore how inbreeding alters these conditions, investigating the effects of sib-mating, lower male-dispersal, oedipal-mating, and the pseudo-hermaphroditism of *Icerya purchasi*, as well as the effect of local resource competition amongst females. Finally, we explore how such genetic and ecological asymmetries may foment conflicts over sexual antagonism between parents and offspring, endosymbionts and their hosts, and maternal-origin and paternal-origin genes.

### Genetic asymmetries

#### The consequences of asymmetric transmission

In most sexual organisms, males and females pass on their maternal-origin and paternal-origin genes with equal frequency. In contrast, haplodiploid organisms are united by the fact that they break this fundamental symmetry, with males exclusively passing on maternal-origin genes (Normark 2006). The best-known form of this is arrhenotoky, whereby males are haploid, produced from unfertilised eggs, and thus carrying only a maternal-origin genome. Consequently, they are constrained to only ever transmit maternal-origin genes, and do so only to daughters. In another form of haplodiploidy, PGE, males are formed from fertilised eggs, and thus initially contain both maternal-origin and paternal-origin genomes. However, either early during development (embryonic PGE) or during spermatogenesis (germline PGE), they eliminate their paternal-genome, and thus their sperm carries only genes of maternal-origin (see Table 1).

**Table 1:**
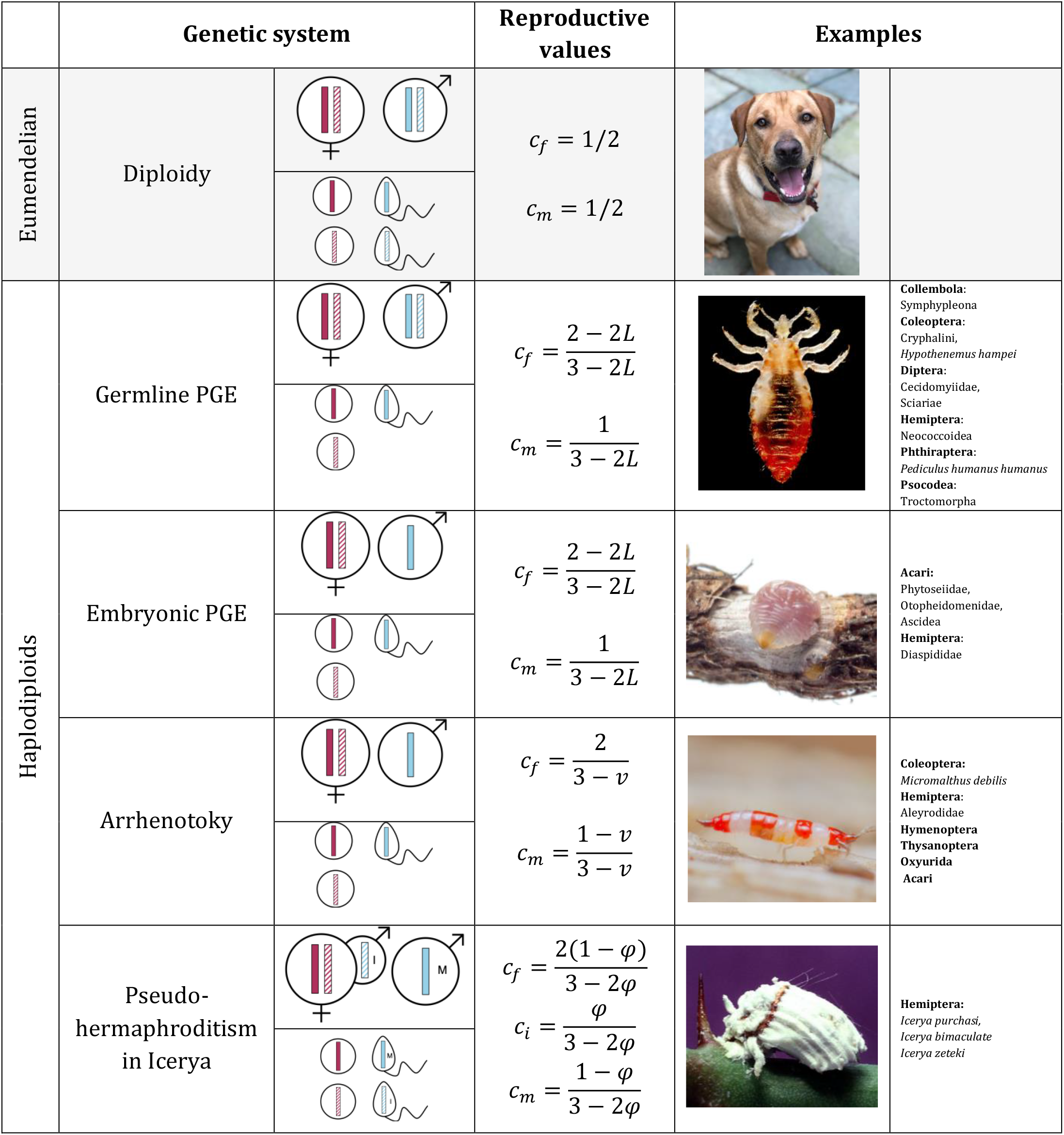
Conceptual description of the different inheritance schemes, and examples of species and groups which fall into these categories, summarised from (Gardner and Ross 2014; de la Filia et al. 2015; Hodson et al. 2017). Solid colours represent maternal-origin genes, and dashed are paternal-origin genes. In PGE systems, *L* is the degree of paternal genome leakage. Under arrhenotoky, *v* is the proportion of oedipal mating. In the Iceryan pseudo-hermaphroditism *φ* is the proportion of eggs fertilised by the infectious male tissue (I), with 1 − *φ* the proportion fertilised by true males (M). Images in order from top to bottom: *Canis familiaris* (Samantha Sturiale), Body louse (public domain), *Pseudaonidia paeoniae* (Matt Bertone), *Haplothrips subtilissimus* (Andy Murray, chaosofdelight.org), *Icerya purchasi* (public domain).

|  | Genetic system |  | Reproductive values | Examples |  |
| --- | --- | --- | --- | --- | --- |
| Eumendelian | Diploidy | | $c_f = 1/2$ | | |
| | | | $c_m = 1/2$ | | |
| Haplodiploids | Germline PGE | | $c_f = \frac{2 - 2L}{3 - 2L}$ | | **Collembola:**<br>Symphypleona<br>**Coleoptera:**<br>Cryphalini,<br>*Hypothenemus hampei*<br>**Diptera:**<br>Cecidomyiidae,<br>Sciaridae<br>**Hemiptera:**<br>Neococcioidea<br>**Phthiraptera:**<br>*Pediculus humanus humanus*<br>**Psocodea:**<br>Troctomorpha |
| | | | $c_m = \frac{1}{3 - 2L}$ | | |
| | Embryonic PGE | | $c_f = \frac{2 - 2L}{3 - 2L}$ | | **Acari:**<br>Phytoseiidae,<br>Otopheidomenidae,<br>Ascidea<br>**Hemiptera:**<br>Diaspididae |
| | | | $c_m = \frac{1}{3 - 2L}$ | | |
| | Arrhenotoky | | $c_f = \frac{2}{3 - v}$ | | **Coleoptera:**<br>*Micromalthus debilis*<br>**Hemiptera:**<br>Aleyrodidae<br>**Hymenoptera**<br>**Thysanoptera**<br>**Oxyurida**<br>**Acari** |
| | | | $c_m = \frac{1 - v}{3 - v}$ | | |
| | Pseudo-hermaphroditism in *Icerya* | | $c_f = \frac{2(1 - \varphi)}{3 - 2\varphi}$ | | **Hemiptera:**<br>*Icerya purchasi*,<br>*Icerya bimaculata*<br>*Icerya zeteki* |
| | | | $c_i = \frac{\varphi}{3 - 2\varphi}$<br>$c_m = \frac{1 - \varphi}{3 - 2\varphi}$ | | |

These distinct transmission genetics alter the relative contributions that females and males make to the ancestry of future generations, i.e. their reproductive values (Price 1970; Taylor 1990; Grafen 2006). Specifically, if we choose a random gene from the distant future and trace it back to the present generation, the probability *c_f_* that it is presently carried by a female defines the class reproductive value of females and the probability *c_m_* = 1 − *c_f_* that it is carried by a male defines the class reproductive value of males, and we find that the ratio of these two reproductive values is given by *c_f_*/*c_m_* = 2(1 − *L*) where *L* is the probability that males transmit their paternal genome, under the assumption of discrete, non-overlapping generations. Under conventional diploidy *L* =1/2 and hence *c_f_*/*c_m_* = 1, i.e. both sexes make an equal genetic contribution to future generations, such that natural selection places equal weight on fitness effects experienced by females and males. In contrast, under haplodiploidy *L* = 0 and hence *c_f_*/*c_m_* = 2, such that females collectively make twice the genetic contribution made by the males, and accordingly natural selection places twice the weight upon fitness effects experienced by females as it does those experienced by males.

If a sexually antagonistic allele confers a marginal fitness benefit *σ* when expressed in females, and a marginal fitness cost *τ* when expressed in males, then the condition for that variant to invade will be *c_f_σ* > *c_m_τ* under outbreeding and in the absence of social interactions between relatives (if the allele were male beneficial then the condition would be *c_f_τ* < *c_m_σ*). A female beneficial allele will therefore invade under haplodiploidy provided 2*σ* > *τ* whilst it will only invade under diploidy provided *σ* > *τ*. Thus, we can see that, relative to eumendelian diploidy, haplodiploidy promotes the invasion of female beneficial alleles (and inhibits the invasion of male beneficial ones).

In some haplodiploid species including mealybugs and body lice, imperfect PGE has been documented, such that in these species males do occasionally transmit paternal-origin genes (de la Filia et al. 2018, 2019). As the amount of male paternal transmission *L* increases, then males obtain an increasing share of the ancestry of future generations, and thus the relative importance of selection on males increases. If paternal transmission were complete (*L* = 1), then all of the future ancestry of the population would reside in males, and thus only fitness effects upon males – either direct or indirect – will have any evolutionary consequences (Gardner and Ross 2011; Yeh and Gardner 2012). In natural populations levels of paternal escape are relatively low (in *Planococcocus citri* the proportion of paternal-transmission was estimated to be between 0.37-3.39%; de la Filia et al. 2019), and thus invasion conditions are essentially identical to strict maternal transmission in males. Nonetheless, slight differences in the degree of leakage between populations, such as the differences documented between ecotypes of *Pediculus humanus* (de la Filia et al. 2018), or potentially experimentally induced paternal-leakage, may allow for effective comparative tests.

#### Asymmetric ploidy and gene effects

Whilst the different haplodiploid systems are united by their common transmission genetics, they often show distinct somatic genetics (Table 1.). These differences in the number of gene copies carried by males and females, and the particular expression patterns of those genes, may alter the relative magnitude of allelic effects in males and females, and thus shape the dynamics of sexual antagonism.

Under arrhenotoky, females carry two genes at each locus, whilst males carry only one. This is conceptually similar to the X chromosomes in an XO system (or an XY system insofar as there is no homologue on the Y) and, as with X chromosomes, it is not necessarily straightforward to compare relative fitness effects across ploidy levels. If an allele’s effect is of similar magnitude in a homozygous and a hemizygous setting, then this will mean that alleles will typically have larger effects on average when expressed in males than in females (Charlesworth et al. 1987; Orr and Otto 1994; Hitchcock and Gardner 2020). For example, if we consider an allele that confers a fitness effect *R* when in hemizygous/homozygous form, then assuming additivity and no inbreeding, a gene expressing this variant strategy will have a marginal fitness effect of *ρ* = *R*/2 when expressed in females, but *ρ* = *R* when expressed in males. Alternatively, if a mutant allele’s effect scales with its absolute rather than relative dosage in the genome (Frank 2003; Gardner 2012; Davies and Gardner 2014), the marginal fitness effects will not systematically differ across the sexes (*ρ* = *R*/2 for males and females). Whilst here we follow the typical assumption of hemizygote/homozygote equivalence, given mechanisms of apparent dosage compensation in some species – such as endopolyploidy in muscle tissues of Hymenoptera (Aron et al. 2005) – then in certain tissues and biological processes it may be more accurate to assume that gene effects scale with absolute copy number (results for these scenarios can be found in the Supplementary Material).

In contrast to arrhenotoky, under PGE, both males and females are initially diploid. If both gene copies within the individual are expressed then, for both males and females, the marginal fitness effects will be *ρ* = *R*/2, as is also the case for eumendelian diploidy. However, among PGE systems there is diversity in the extent of somatic paternal genome expression. This may occur either because the whole or part of the paternal genome is eliminated early in development (embryonic PGE), such that somatic tissues are actually haploid, or because the paternal-genome is silenced, such that certain tissues are functionally haploid (Burt and Trivers 2006; de la Filia et al. 2021). If a locus is exclusively maternally expressed, then marginal fitness effects are identical to those given for arrhenotoky. Thus, depending on species, tissue, and locus, we expect to observe a continuum between these two scenarios (Figure 1). For simplicity, we henceforth assume that both gene copies are fully expressed under PGE, a scenario that captures autosomal expression in several PGE clades including springtails, lice and fungus gnat and gall midge flies (de la Filia et al. 2015, Table 1). It also captures the evolution of a subset of genes and tissues in species where paternal genome is silenced, such as mealybugs, as silencing appears to be incomplete (de la Filia et al. 2021). In contrast species with germline PGE (Table 1) are equivalent to arrhenotokous species as males become fully haploid early in development.

**Figure 1:**
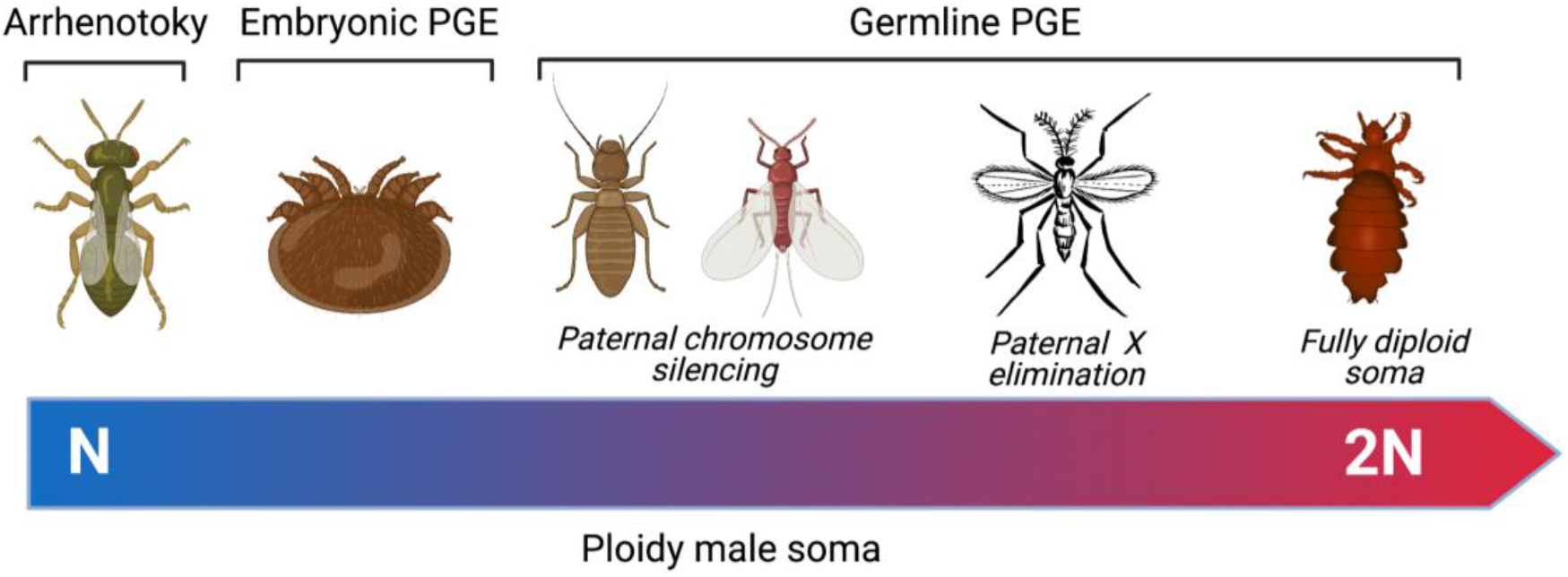
The continuum of male gene expression and thus effective ploidy level found across haplodiploid groups with representative taxa illustrated, from left to right, *Nasonia vitripennis,* Varoa mite (*Varoa destructor*), *Liposcelis* booklice, citrus mealybug male (*Planococcus citri*), Hessian fly (*Mayetiola destructor*) and head louse (*Pediculus humanus capitis*)

A further factor that may modulate the relative scaling of gene effects across sexes is dominance (Rice 1984; Fry 2010; Patten 2019). Relaxing the assumption about additivity, then under diploidy and PGE a gene’s marginal fitness effect will be *ρ* = *h_m_R* in males and *ρ* = *h_f_R* in females, but under arrhenotoky will be *ρ* = *R* in males and *ρ* = *h_f_R* in females. With no sex differences in dominance then this will have no systematic effect on PGE and diploidy, as it will impact both sexes equally. In contrast, under arrhenotoky, as dominance only impacts upon diploid females, then it will scale fitness effects between males and females. Further complications arise if there are reversals of dominance, i.e. the beneficial and deleterious effects of sexually antagonistic mutations are expected to have reversed dominance coefficients (Fry 2010; Patten 2019). For simplicity, we restrict our attention here mostly to the additive case, however full results for non-additive scenarios can be found in the Supplementary Material.

Integrating the weightings from transmission with the marginal fitness effects, we find that, under outbreeding, the condition for a female beneficial allele to invade will be *S* > *T* under arrhenotoky (and diploidy), but 2*S* > *T* under male PGE, where *S* and *T* are the fitness costs and benefits of the allele to hemizygous/homozygous individuals (see Table 2.). For a male beneficial allele, the conditions will be *S* > *T* under arrhenotoky (and diploidy), but *S* > 2*T* under male PGE. We can see that with arrhenotoky, the two-fold weighting placed on females is cancelled out by the two-fold larger fitness effects in males, whilst under male PGE, where marginal fitness effects are not systematically different across sexes, this cancellation does not occur. Thus, we would generally expect relative feminisation of the genome in PGE species as compared to arrhenotokous ones, as invasion conditions for female-beneficial alleles are less stringent, and those male-beneficial alleles are more stringent (see Figure 2).

**Table 2:**
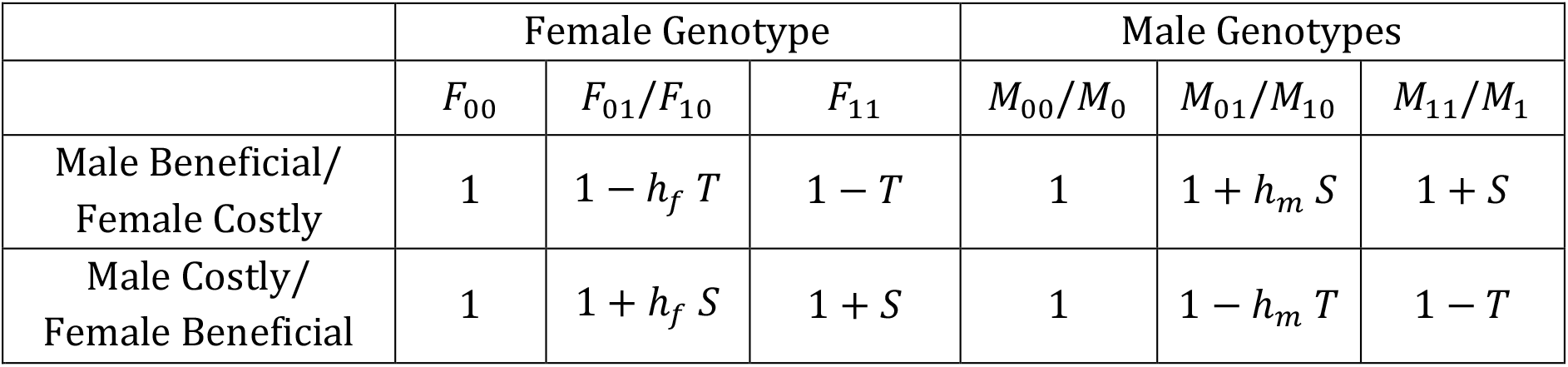
Fitness scheme for invasion analysis.

**Figure 2:**
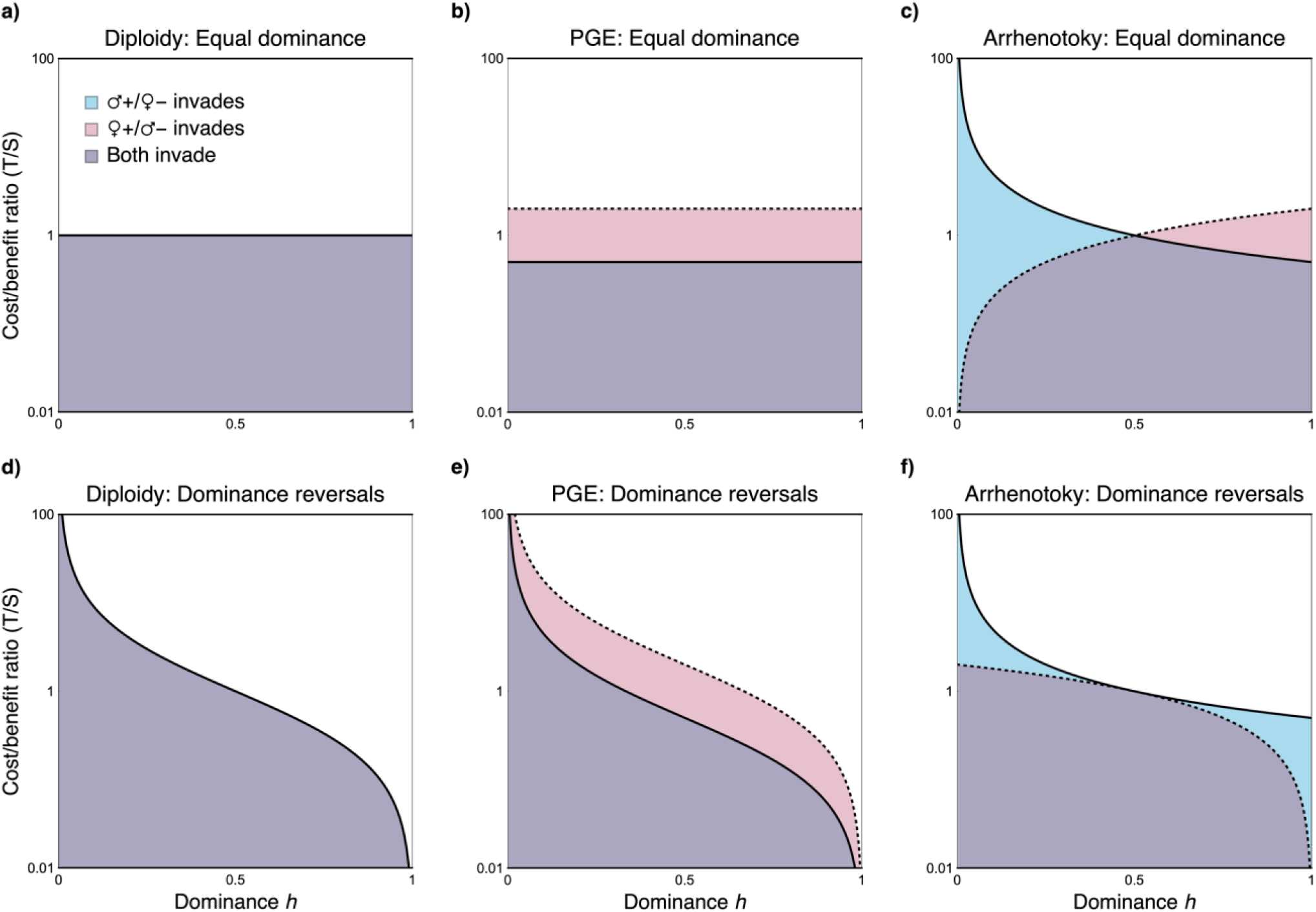
Female beneficial alleles invade more readily under germline PGE than they do under arrhenotoky. The invasion space for sexually antagonistic mutations with a given cost/benefit ratio (*T*/*S*) under different inheritance schemes and assumptions about dominance (*h*), with male beneficial alleles invading beneath the solid line, and female beneficial alleles beneath the dotted line. In the equal dominance scenarios (a-c): *h* = *h_f_* = *h_m_* for both male and female beneficial alleles. In the reversals of dominance scenarios (d-f): *h* = *h_f_* = 1 − *h_m_* for the male beneficial scenario, and *h* = 1 − *h_f_* = *h_m_* for the female beneficial scenario.

### Ecological asymmetries

#### Sib-mating and ecological asymmetries between the sexes

The above results apply to outbreeding populations with no social interactions between relatives, and therefore it is only the direct fitness effects of alleles that required consideration. But many haplodiploid species diverge from this, with mating schemes and life-cycles that result in chronic inbreeding (Hamilton 1967, 1978, 1993). These population structures may alter the relatedness within and between individuals, as well as the intensity with which males and females compete with relatives, potentially generating indirect fitness effects of sexually antagonistic alleles upon social partners. Such factors have long been recognised in sex allocation research to alter the relative value of sons and daughters (Taylor 1981; Frank 1986b; Nagelkerke and Sabelis 1996; West 2009), and thus may be expected to play a similar role with regards to sexual antagonism.

We investigate how inbreeding may modulate sexual antagonism by modelling a population of monogamous females, in which a proportion *s* of females in the brood mate with their sibs, whilst a proportion 1 − *s* mate with males from the population at large (Figure 3). Introducing sib-mating has multiple distinct effects upon sexual antagonism. The first is that sib-mating inflates the consanguinity of an individual to themselves, i.e. their inbredness (*sensu* Frank 1986a), which has a feminisation promoting effect under arrhenotoky – as a gene copy will have indirect fitness effects upon the other identical by descent gene copy in females, but not in males, which are haploid (Tazzyman and Abbott 2015; Hitchcock and Gardner 2020) – but not for PGE or diploidy, where gene copies in both males and females experience these within individual indirect fitness effects. Secondly, sib-mating increases the probability that males will compete with brothers for mates, discounting the inclusive-fitness benefits of male-beneficial alleles to their male carriers, and mollifying the inclusive-fitness costs of male-deleterious alleles. Thirdly, the direct fitness effects of alleles upon their female carriers will have indirect fitness effects upon their carriers’ mates. If females sib-mate, then female-beneficial alleles will generate indirect benefits for their brothers, and female-deleterious alleles will impose indirect costs. All three of these effects have parallels in sex allocation, with increased sib-mating increasing the relatedness of a female to her daughters but not her sons under arrhenotoky, increased competition between brothers decreasing the genetic returns on males (i.e. local mate competition; Hamilton 1967), and increased sib-mating meaning that increased investment into daughters will increase the fitness of sons, either through extra mating opportunities, or through higher quality mates (Taylor 1981; Frank 1986b; West 2009).

**Figure 3.**
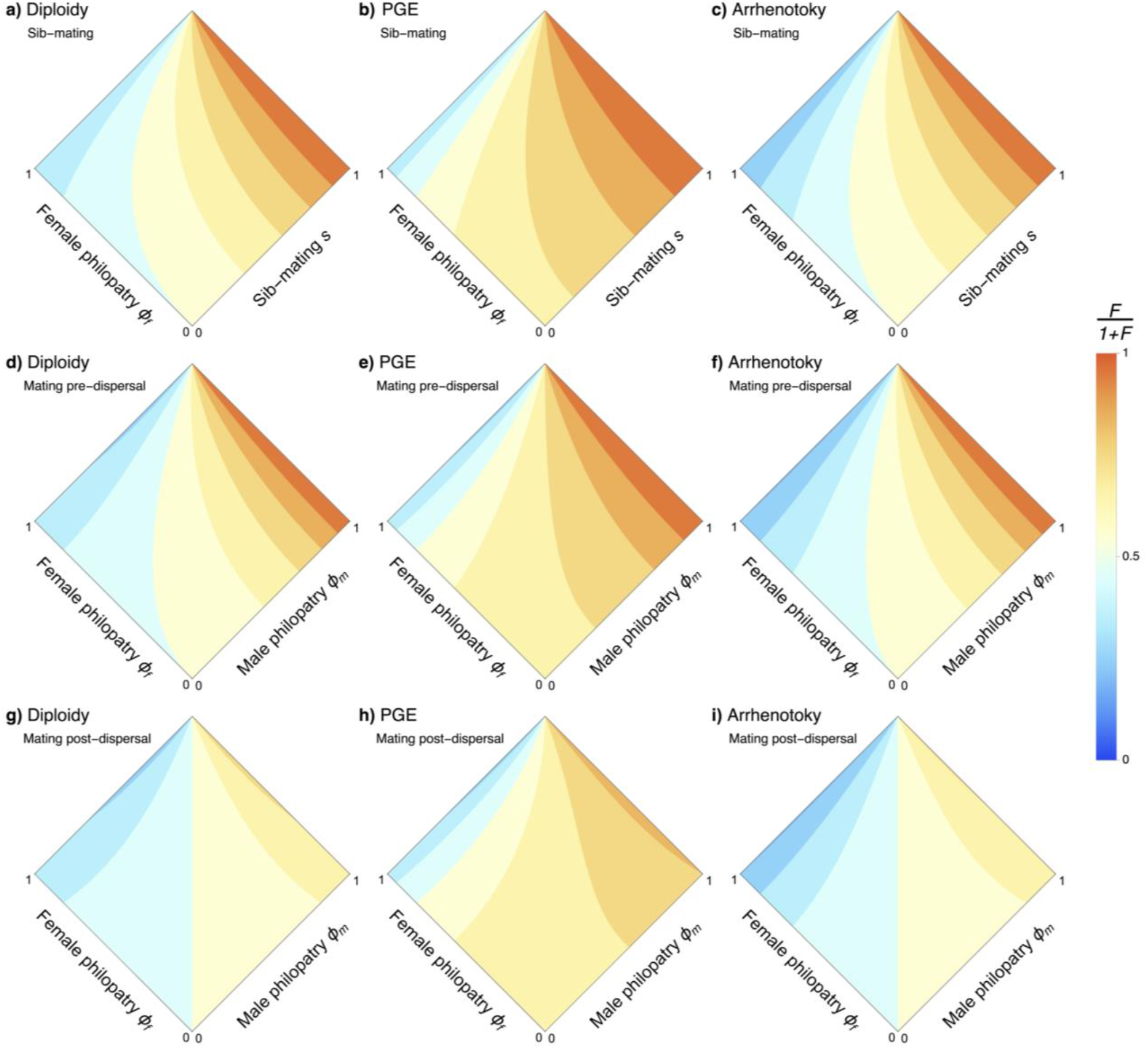
Mating ecology and dispersal modulate the degree of feminisation. Here the degree of feminisation, *F*/(1 + *F*), is plotted as a function of either the amount of male and female philopatry, or the amount of female philopatry and the proportion of sib-mating, under three inheritance systems (diploidy, germline PGE, and arrhenotoky), and for three mating ecologies (sib-mating (a-c), viscous population with mating pre-female dispersal (d-f), and viscous population with mating post-female dispersal (g-i)).When *F*/(1 + *F*) > 0.5, then feminisation is expected, and when *F*/(1 + *F*) < 0.5 masculinsation is expected.

Collecting these effects, we may write the condition for a female beneficial to invade in the form of a potential for feminisation *F*, with a female beneficial allele invading provided *F* < *T*/*S*. If *F* > 1, then feminisation is expected, whilst if *F* < 1 then masculination is expected. Assuming additivity and weak selection, we find that under arrhenotoky and diploidy *F* = 1/(1 − *s*), and under male PGE *F* = (4 − s)/(2 (1 − s)) (results for dominance and stronger selection can be found in the Supplementary Material). Thus, we find that, across all these genetic systems, increased sib-mating promotes feminisation, with the effect being strongest under PGE (see Figure 3).

So far, we have assumed that females compete globally, however, many haplodiploid species have more generally viscous populations in which females may also disperse short distances – if at all. For instance, in the date stone beetle, females may start their own families within the seed in which they were born (Spennemann 2019). Similarly, in many mealybugs, females crawl relatively small distances away from their natal patch (Varndell and Godfray 1996; Ross et al. 2010a). In these species females may compete with sisters for breeding spots, just as their brothers competed with each other for mates, i.e. local resource competition (Clark 1978). Incorporating these factors yields two further consequences for sexual antagonism. Firstly, with limited female dispersal, direct fitness benefits to females incur indirect fitness costs to their sisters by depriving them of breeding spots, just as obtained for local mate competition in males. Secondly, whilst a fit female confers indirect fitness benefits upon brothers with whom she mates, she may also incur indirect fitness costs by competing with her brothers’ mates, and thereby indirectly depriving her brothers of reproductive success. With increasing local resource competition, the invasion condition becomes less stringent for male-beneficial alleles and more stringent for female-beneficial alleles. The dual effects of sib-mating and limited female dispersal can be seen in Figure 3, with full analytical results given in the Supplementary Material.

#### Alternative life-cycles and modes of inbreeding

Above we considered one particular inbreeding scenario, in which a fixed proportion of matings are reserved for siblings. However, the specific mechanism by which inbreeding occurs may also modulate sexual antagonism, as different mating schemes and life-cycles will differ in how relatedness builds up, and how intensely males and females compete with relatives. To investigate this, we contrast the above model with an alternative involving a patch structured population in which the degree of inbreeding is modulated by the extent of dispersal (Wright 1931), whereby males remain on their natal patch with probability *ϕ_m_*, and females with probability *ϕ_f_*. We consider two variants, the first in which mating occurs before female dispersal (male dispersal → mating → female dispersal, DMD), and a second in which mating occurs after female dispersal (male dispersal → female dispersal → mating, DDM), the latter of which has been recently investigated by Flintham et al. (2021) for sexual antagonism in relation to diploidy and X-linkage. Comparing these results, we obtain a ranking of highest potential for feminisation under sib-mating, followed by DMD, and finally DDM (Figure 3, see Supplementary Material for details). The sib-mating and DMD scenarios are very similar, except that brothers are more likely to compete for mating opportunities in the former scenario, promoting feminisation, analogous to the difference between fixed self-fertilization and mass-action selfing models of hermaphroditic plants (Jordan and Connallon 2014). Compared to DDM, both sib-mating and DMD scenarios yield a higher potential for feminisation, as they involve both higher rates of consanguineous mating and also sisters conferring fitness benefits upon related mating partners, an effect that is exactly cancelled under DDM by increased competition between females and their brothers’ mates. Thus, different mating ecologies and life-cycle structures yield different patterns of feminisation.

Alongside the generic demographies discussed above, haplodiploids present a striking variety of unusual lifecycles and modes of inbreeding. For illustration, we consider two scenarios in detail, both of which involve females effectively engaging in ‘selfing’. First, oedipal mating (Table 1.) occurs because a virgin female may produce an exclusively male brood with which she then mates, a reproductive strategy observed in groups including: mites (McCulloch and Owen 2012; Tuan et al. 2016) beetles (Entwistle 1964; Jordal et al. 2001), parasitoid wasps (Browne 1922; Schneider et al. 2002), pinworms (Adamson and Ludwig 1993), and thrips (Ding et al. 2018). Second, in the scale insect *Icerya purchasi*, selfing is understood to occur as a consequence of a diploid female containing a transovarially transmitted haploid spermatogenic cell lineage that may fertilise her eggs (Royer 1975; Normark 2009; Ross et al. 2010b; Mongue et al. 2020). Whilst these two systems are very different in their biological details, in both cases we find that higher rates of ‘selfing’ increases the potential for feminisation, and do so in a fashion that is qualitatively very similar to sib-mating (see Supplementary Material for full details).

### Conflicts over sexual antagonism

#### Parent-offspring conflict over sexually antagonistic traits

In the foregoing, we have assumed that the sexually antagonistic traits of interest are under the sole control of the individuals in which they are expressed. However, the traits of an individual may also be influenced by social partners. In particular, parents may play an important role in shaping the traits of their offspring, whether it be through the material constitution of the zygote, the environment in which those offspring develop, or through the care that those parents provide (Mousseau and Dingle 1991; Mousseau and Fox 1998; Crean and Bonduriansky 2014). If the traits that they influence are sexually antagonistic, then parents may face a trade-off between crafting superior daughters versus superior sons. Moreover, if parents place different values upon males and females as compared to their offspring, then this may lead to parent-offspring conflict (*sensu* Trivers (1974)) with respect to sexually antagonistic traits. Furthermore, if mothers and fathers also differ in their relative valuations of sons and daughters then this may lead to sexual conflict (*sensu* Trivers (1972)) with respect to sexually antagonistic traits.

Focusing our attention first on genes acting through mothers, if we consider the invasion of an allele which increases the fitness of her daughters, but decreases the fitness of her sons, then for diploidy the potential for feminisation may be expressed as *F* = (1 + *s*)/(1 − *s*). When there is no sib-mating, then this is equivalent to that for offspring, a result previously found when considering organisms with a dominant haploid phase (Patten and Haig 2009). However, under sib-mating the interests of mothers and offspring diverge, with mothers favouring a greater female bias than their offspring (Figure 4a). This parallels a previous effect found in relation to sex allocation, whereby offspring typically favour less extreme sex ratio deviations than their parents (Trivers 1974; Werren and Hatcher 2000; Pen 2006), on account of parents being favoured to maximize the success of the entire brood whereas each individual values itself more than its siblings (although see Pen (2006) for situations where this pattern may be reversed). For arrhenotoky and PGE, *F* = [3 − (1 − *s*)^2^]/[(1 − *s*)(2 − *s*)]. Thus, for arrhenotoky, the situation is similar to diploidy, with mothers and offspring in agreement under random mating, but with mothers favouring a greater female bias when there is sib-mating (Figure 4c). For PGE, however, when there is no sib-mating then offspring favour more female bias than their mothers, as females are twice as valuable as males from the perspective of the offspring, whilst sons and daughters are equally valuable from their mothers’ perspective. However, this situation reverses as sib-mating increases, with mothers once again favouring more female-biased trait values than their offspring (Figure 4b).

**Figure 4:**
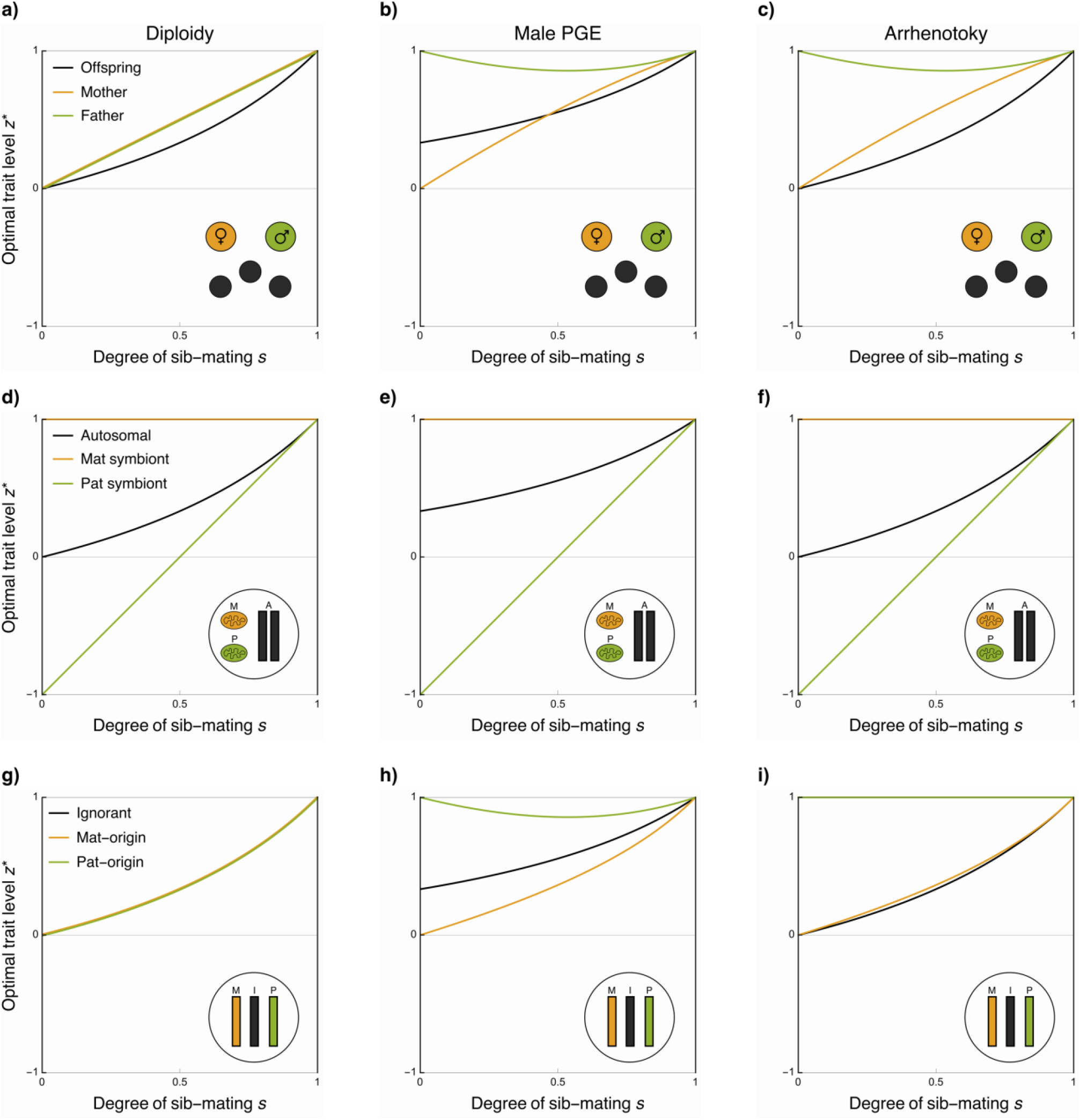
Conflicts within and between individuals over sexually antagonistic traits, across different genetic systems. The optimal level of a sexually antagonistic trait *z* under diploidy, germline PGE, and arrhenotoky when control of that trait is assigned to: offspring, mothers, and fathers (a-c); autosomal genes, matrilineal cytoplasmic genes, and patrilineal cytoplasmic genes (d-f); ignorant genes, maternal-origin genes, and paternal-origin genes (g-i). In these examples, fitness is a gaussian distributed trait with an optimum of 1 for females and −1 for males, with equal variance.

Considering instead a sexually antagonistic allele that acts through fathers, we find that for diploidy the potential for feminisation is the same as for mothers, *F* = (1 + *s*)/(1 − *s*), and with both parents favouring a more female biased trait value than offspring. For arrhenotoky, however, fathers favour a far more feminised trait value than either offspring or mothers, *F* = [3 + (1 − *s*)^2^]/[(1 − *s*)*s*], as they only contribute genetically to their daughters in the brood, similarly to how fathers favour an exclusively female brood under outbreeding (Hamilton 1967). Nonetheless, with increased sib-mating they are increasingly related to their mates’ sons, and thus place value on their fitness too, but with further sib-mating this is counteracted by the effects of increased local mate competition, once again favouring feminisation (Fig 4c). PGE yields a qualitatively similar outcome; however, as a male’s paternal-origin genome is passed to neither sons nor daughters directly, then fathers are not as highly related to their daughters as compared with arrhenotoky, and they therefore favour slightly less feminisation (Fig 3b), with the potential for feminisation *F* = [6 − (1 − *s*)(2 − *s*)(1 + *s*)]/[(1 − *s*)(3 − *s*)*s*].

#### Endosymbionts, mitochondria, and germline restricted chromosomes

Thus far we have largely treated the genome as though it is a unified entity. However, even though different genes may reside within the same body, they may nevertheless have distinct inclusive-fitness interests (Hamilton 1967; Burt and Trivers 2006; Gardner and Úbeda 2017), and thus come into conflict over the trade-offs imposed by sexual antagonism. This is particularly relevant for haplodiploids as many contain endosymbionts which have different transmission modes to autosomal genes (Buchner 1965; Normark 2004a; Ross et al. 2012; Perlmutter and Bordenstein 2020), and thus may place different valuations upon males and females (Hurst 1991; Frank and Hurst 1996). Similarly, particular species also contain further unusual genomic features, such the matrilineally-inherited germline restricted E chromosomes found in gall midges (Harris et al. 2003; Normark 2004a; Hodson and Ross 2021).

For those endosymbionts and chromosomes that are strictly matrilineally inherited, they will place no direct value upon the fitness of males, bringing them into conflict with the rest of the genome (Wade 2014; Hurst and Frost 2015). These elements may also therefore provide a rich source of evidence for the “Mother’s Curse” hypothesis, i.e. that mitochondria accumulate mutations which are deleterious for males (Gemmell et al. 2004). Under full outbreeding this conflict is at its most intense, but with increasing amounts of sib-mating the autosomes becomes increasingly female biased too, aligning the interests of these two sets of genes, and thus reducing the extent of the conflict. This also applies to patrilineally-inherited symbionts, which although much rarer than matrilineally inherited counterparts have been documented in a variety of species including aphids (Moran and Dunbar 2006), mosquitos (Damiani et al. 2008), leafhoppers (Watanabe et al. 2014), termites (Korb and Aanen 2003), and tsetse flies (De Vooght et al. 2015). With full outbreeding, paternally-inherited genes place no value on females, but as inbreeding increases then they place an increasing value on the fitness of females, mollifying the conflict between them, autosomal, and maternal-inherited genes, as shown in Figure 4d-f.

#### Parent-of-origin specific gene expression

Finally, a further intragenomic conflict that may emerge over sexual antagonism is that between maternal-origin and paternal-origin genes (Haig 2002). The asymmetric transmission genetics that defines haplodiploidy may subsequently generate differences between maternal-origin and paternal-origin genes in how they value males and females, and also their relatedness to the males and females with whom they interact (Haig 1992; Queller and Strassmann 2002; Queller 2003; Wild and West 2009; Rautiala and Gardner 2016; Marshall et al. 2020).

In the simplest case, with full outbreeding, we find that if a gene is of maternal-origin it places equal value upon males and females, under diploidy, arrhenotoky and PGE. Conversely, if it is of paternal-origin then it places equal value upon males and females under diploidy, but places no value upon males under the haplodiploid systems, as it is never transmitted by males under PGE, and is absent from males under arrhenotoky. Focussing on PGE, we can explore how, depending on which gene copy controls the trait, the potential for feminisation may change. This is particularly relevant as the extent of expression in males from the maternal-origin and paternal-origin copies may vary across loci, tissues, and species (Burt and Trivers 2006; Gardner and Ross 2014; de la Filia et al. 2021). Allowing for a proportion *y* of a locus’s expression in a male to come from the paternal-origin copy, and a proportion 1 − *y* to come from the maternal-origin copy, we find that the potential for feminisation is *F* = 1/(1 − *y*). Thus, when maternal-origin genes control the trait in males (*y* = 0), then *F* = 1, equivalent to the arrhenotokous case, whist when expression is exclusively from the paternal-origin copy (*y* = 1), then *F* = ∞, i.e. female beneficial alleles will always invade, regardless of the cost they impose upon males, analogous to how paternal-origin genes may favour male suicide when there is competition between male and female siblings (Ross et al. 2011b).

As the rate of sib-mating increases, the intragenomic conflicts become more complex. We now explore the effects of parent-of-origin specific gene expression in both males and females. Allow for a proportion *y* of a locus’s expression in males to come from their paternal-origin copy and a proportion 1 − *y* from their maternal-origin copy, and allowing a proportion *x* of that locus’s gene expression in females to come from their maternal-origin copy, and proportion 1 − *x* from their paternal-origin copy. Then we find the degree of feminisation under PGE becomes:

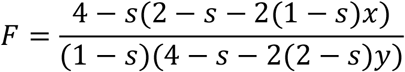

With the results for arrhenotoky generated by setting *y* = 0. We can see that assigning full control to maternal-origin copy in both sexes (*x* = 1, *y* = 0), conditions simplify to *F* = [(2 + *s*)(2 − *s*)]/[(4 − *s*)(1 − *s*)], which is a monotonically increasing function of *s*, i.e the degree of feminisation always increases with increasing *s*. In contrast, if we assign full control to the paternal-origin genes (*x* = 0, *y* = 1), then *F* = [3 + (1 − *s*)^2^]/ [(1 − *s*)*s*]. When *s* = 0, then the paternal-origin copy is unrelated to the other gene copy in a male, and thus places no value on male fitness. As *s* increases then the value that a paternal-origin gene places on males increases too, as that gene copy is related to the other gene copy it resides in a male with. However, with further increases in *s*, this is countered both by the increasing competition between related males, and also the indirect effects from related females.

Previously, intragenomic conflict between maternal-origin and paternal-origin genes has been suggested to drive the evolution of genomic imprinting at such loci, i.e. the expression of one parental copy, and the silencing of the other parental copy. This results from an escalating conflict over joint expression levels, which ultimately results in the gene copy that favours lower expression levels becoming silenced, whilst the one that favours higher expression levels is expressed at its optimum level, a process termed the “loudest voice prevails” principle (Haig 1996). If we apply the logic of this principle to conflicts over sexually antagonistic traits then, under PGE, we may expect paternal-origin genes to be expressed for female-beneficial trait promoters and male-beneficial trait inhibitors, whilst we would expect maternal-origin genes to be expressed for male-beneficial trait promoters, and female-beneficial trait inhibitors. These predictions are distinct from those made under other theories of sexually antagonistic genomic imprinting (Day and Bonduriansky 2004), whereby we would expect paternal-origin genes to be expressed in males, and maternal-origin genes to be expressed in females.

## Discussion

Haplodiploid species account for a large minority of all animal species (Normark 2003, 2006; Bachtrog et al. 2014; de la Filia et al. 2015), with many striking examples of sexual dimorphism (see Table 1, Figure 5). Our analyses here have shown how some of the unusual genetic and ecological asymmetries that define these groups are expected to modulate the outcome of sexual antagonism. We find that PGE systems are broadly more favourable to the invasion of female-beneficial alleles (and less favourable to their male-beneficial counterparts) as compared to arrhenotoky and diploidy, that the chronic inbreeding associated with many of these species promotes feminisation, and that both of these factors may foment conflicts between different parties over sexually antagonistic traits. We have also considered how some of the diversity of genetics and life-history exhibited by haplodiploids may further modify these results, generating variation in the fate of sexually antagonistic alleles across loci, tissues, and species.

**Figure 5:**
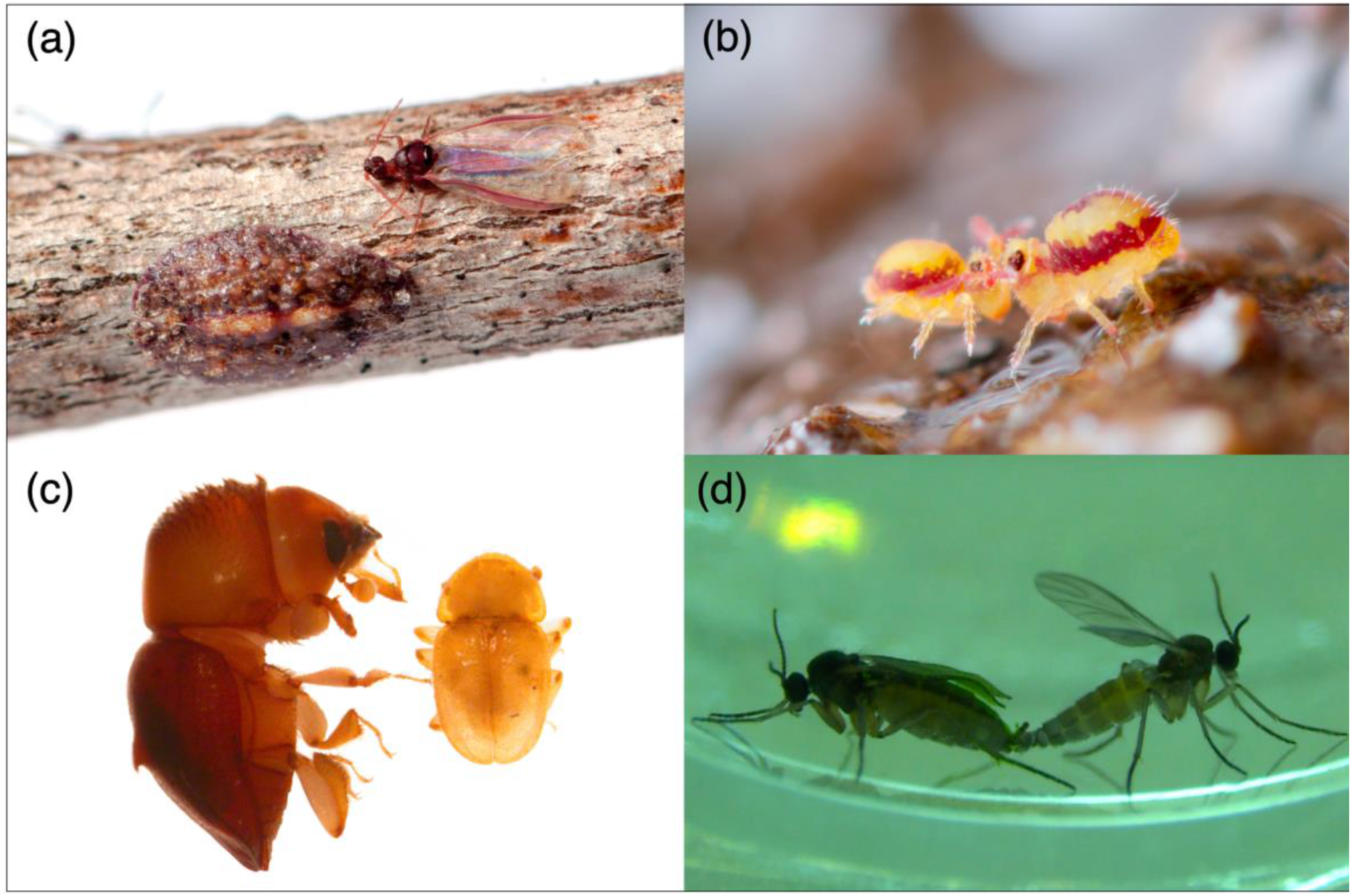
Examples of sexual dimorphism in haplodiploid species. a) Soft scale insects (*Pulvinaria acericola*), female on the left, male on the right (credit: Matt Bertone). b) Globular springtail (*Sminthurides malmgreni*), male on the left, female on the right (credit: Andy Murray, chaosofdelight.org). c) Ambrosia beetle (*Diuncus sp.*) female on the left, male on the right (credit: Jiri Hulcr). d) Fungus gnats (*Bradysia coprophila*), female left, male right (credit: Robert Baird).

Whilst our analysis indicates that these groups may provide a particularly rich set of comparative tests for how ecology and genetics modulate sexual antagonism, relatively little work has been carried out to investigate this (although see Table 2 for a summary). One of the reasons for this is that the within-genome comparisons often used to study sexual antagonism have been considered impossible for the many haplodiploid species that lack sex chromosomes. However, this overlooks the exceptions that provide excellent opportunities for testing theory. For instance, sciarid flies not only have male PGE, but also an XO sex chromosome system (Metz 1938; Rieffel and Crouse 1966), thus allowing a within-organism comparison of these inheritance systems in relation to sexual antagonism. This is also true of some other groups with germline PGE such as gall midges and globular springtails (Gallun and Hatchett 1969; White 1977; Dallai 2000; Anderson et al. 2020). In these groups we may expect female-beneficial variants to be enriched on the autosomes, whilst male-beneficial ones would be expected to be overrepresented on the sex chromosomes, regardless of assumptions about dominance, making this a more straightforward prediction than between autosomes and sex chromosomes in conventional eumendelian systems (Rice 1984; Patten 2019). In addition to X autosome comparisons, some of these groups contain further genomic elements such as germline-restricted chromosomes that are maternally-inherited in gall midges and show paternally-biased inheritance in sciarid flies (Hodson and Ross 2021; Hodson et al. 2021), enabling further within-genome comparisons.

Similarly, whilst it has been suggested that the X chromosome should be relatively enriched for sexually antagonistic polymorphisms in eumendelian systems as compared to the autosomes (Rice 1984), again this depends on assumptions about dominance (Fry 2010; Ruzicka and Connallon 2020). We find here that the same is true of comparisons between PGE and X chromosomes/arrhenotoky, with arrhenotokous organisms ones having a higher potential for polymorphism under parallel dominance, but a smaller space for polymorphisms under dominance reversals (see Supplementary Material). Additionally, such sexually antagonistic polymorphisms may be easier to detect in some haplodiploid species as compared to eumendelian ones, because the asymmetric transmission genetics means that allele frequency differences that build up between the sexes in one generation, will carry over to the next (Crow and Kimura 1970; Ruzicka and Connallon 2020).

Additionally, we find that the chronic inbreeding exhibited by many haplodiploids typically promotes feminisation. This meshes with the increasing interest in the role of demography and ecology in modulating sexual antagonism (Albert and Otto 2005; Arnqvist 2011; Harts et al. 2014; Tazzyman and Abbott 2015; Connallon et al. 2019; de Vries and Caswell 2019; Hitchcock and Gardner 2020). In particular, Flintham et al. (2021) have recently shown how, in viscous populations, sex-biased dispersal may skew sexual antagonism under diploidy and X-linkage towards the sex that competes less intensely with relatives. Here we recover that same pattern, but also find that other mating schemes that characterise haplodiploid groups can involve an additional feminising effect, as females may confer fitness benefits upon their mates. Alongside comparisons between populations and species, one method of testing such predictions would be through the use of experimental evolution. For example, Rodrigues et al. (2021) evolved populations of the spider mite *Tetranychus urticae* under various dispersal regimes in order to investigate the evolution of sex allocation; those demographies predicted to lead to greater female bias in the sex ratio would also be expected to promote female bias in relation to sexual antagonism. Thus, under these conditions, we may expect to see either increased fixation of female-beneficial sexually antagonistic alleles and/or phenotypes moving toward the female optimum. Reinvestigation of these evolved lines or new experiments with similar design would enable testing of predictions emerging from our analysis.

Furthermore, we have shown how population structure and transmission asymmetries may foment conflicts between different genetic parties over sexually antagonistic traits. In particular, we identify potential for conflict between parents and offspring. Whilst there has been similar work considering the differing interests between parents and offspring with regards to sex allocation (Trivers 1974; Werren and Hatcher 2000; Pen 2006), sexual antagonism provides a further arena for such conflicts of interest. Whilst parent-offspring conflict emerges across all of our genetic systems under sib-mating, species with PGE provide a particularly interesting set of systems within which to investigate this phenomenon as, even under full outbreeding, mothers, fathers, and offspring all favour different trade-offs. Thus, depending on who controls the trait, we may expect different patterns of masculinisation vs feminisation. Comparisons between sperm-derived versus egg-derived products, and between those to genes expressed after the maternal-to-zygotic transition, may help reveal such conflicts over development. A further, particularly interesting case to investigate the logic of such conflicts is with the bacteriome of the armoured scale insects. These are pentaploid tissues containing two complete copies of the mother’s genome and a copy of the paternal-origin genome (Normark 2004b). Thus whilst not identical to the parents interests, the bacteriome nonetheless might be expected to have more similar genetic interests to the mother than the offspring it resides within, and thus the interface between them provides a within-individual arena for this parent-offspring conflict.

We have focused here on cases where there are only two classes of individual: males and females. However, many of the better known haplodiploid species – most notably the eusocial Hymentoptera – exhibit not just sex-structure, but also caste-structure. For instance, in the eusocial bees, wasps, and ants, in addition to reproductive females (queens) and reproductive males (drones), there is also an additional female neuter class (workers) who are morphologically, physiologically, and behaviourally distinct from the queen. Whilst the addition of caste structure on its own is not expected to modulate sexual antagonism per se, i.e. trade-offs between queens and reproductive males, if the trade-off occurs through female workers and reproductive males then results would be expected to diverge, as phenotypic effects that manifest in females would only have indirect effects through their effects on the reproductive females. Moreover, with more than two castes there is the possibility for more complex trade-offs operating across multiple classes, such as between workers and queens, workers and males, and three-way trade-offs; such trade-offs have previously been referred to in terms of ‘intralocus caste antagonism’ (Holman 2014; Pennell et al. 2018). A similar complexity occurs when males exhibit polyphenisms, for instance in fig wasps between winged and non-winged male forms (Hamilton 1979; Cook et al. 1997). Such male dimorphism can be extreme, not only concerning the presence/absence of wings, but also with respect to other aspects of morphology and behaviour. If a sexually antagonistic allele affects these morphs differently, then outcomes will be more complex than those emerging from our analysis, depending on the relative fraction of male dispersers. Similarly to caste structure, this may lead to trade-offs amongst these male morphs, previously termed ‘intralocus tactical evolution’ (Morris et al. 2013).

Our predictions have been derived under the assumption of non-overlapping generations, yet age-structure may also have an important modulating effect on sexual antagonism (de Vries and Caswell 2019; Hitchcock and Gardner 2020). This may be important for two reasons. Firstly, sex-specific age-structure may disturb the reproductive values of males and females away from the ratios given here (Grafen 2014; Hitchcock and Gardner 2020). This may be because there are sex-differences in mortality and fecundity, such as in the citrus mealybug (*Planoccocus citri*) where males live up to only 3 days post eclosion whilst females can live several weeks (Nelson‐Rees 1960; Ross et al. 2011a), or because of other factors which can generate more cryptic age-structure such as partial bivoltinism (Seger 1983; Grafen 1986), sperm storage, or worker reproduction (Benford 1978; Charnov 1978; Alpedrinha et al. 2013). Secondly, population viscosity may generate competition between parents and offspring (Irwin and Taylor 2001; Ronce and Promislow 2010). Coupled with other aspects of sex-biased demography, such as sex-biased dispersal (Johnstone and Cant 2008, 2010), then this may reduce the magnitude of costs/benefits to one sex more than the other, and thus bias the outcome of sexual antagonism toward one sex. An example relevant to this is the aforementioned date-stone beetle where a single female may spawn up to five generations within a single drupe over the spring and summer (Spennemann 2019), thus generating potentially strong inter- and intra-generational kin competition.

Finally, we have considered mating to be the only social interaction between males and females. Yet invasion conditions for sexually antagonistic alleles are liable to be modulated by more extensive and complex intersexual interactions. For instance, intrabrood competition may result in male-beneficial alleles decreasing the fitness of females both through the direct effect of those alleles being expressed by females, but also through those females being outcompeted by their brothers (and vice versa, for female-beneficial alleles). The extent of such competition will vary with ecological context. For instance, bark beetles are understood to experience intense sib-competition, whilst phloem feeders are less likely to do so (Normark 2004a, 2006). Intense intrabrood competition is also an ecology well-suited to the evolution of cytoplasmic male killing (Hurst 1991; Hamilton 1993; Normark 2004a). Moreover, we have assumed that there is an asymmetry in that female-beneficial variants improve the likelihood of a mating pair winning a breeding opportunity (as it is competitiveness of females that determines this), whilst male-beneficial variants have no such effect. Whilst this does adequately capture the ecology of many haplodiploid species, there are scenarios in which this assumption need not hold. For instance, males may have beneficial fitness effects upon their mates if there is paternal care, as in the case of the mud daubers (Brockmann 1980; Bragato Bergamaschi et al. 2015) and the solitary apid bee, *Ceratina nigrolabiata* (Mikát et al. 2019), or if sperm is a limiting factor on the rate of reproduction. Alternatively, males may also have deleterious fitness effects if they exhibit harming traits, such as the traumatic insemination observed in some groups of pinworms (Adamson 1989).

In conclusion, we have explored how genetic and ecological asymmetries that characterise haplodiploid groups are expected to modulate sexual antagonism, and how these may in turn foment conflicts both between and within individuals over such traits. Exploring the consequences of these unusual genetic systems and life-cycles has previously offered rich insights into sex allocation (Charnov 1982; West 2009), and thus leveraging the natural diversity within these groups may also deepen our understanding of sexual antagonism and the evolution of sexual dimorphism. Gene expression studies increasingly look for sex biased gene expression in such non-model and non-eumendelian species, and our predictions will facilitate interpretation of these data, as well as identifying where future research effort may be most fruitfully focused. Finally, many of the species that reproduce through arrhenotoky or PGE are pests and parasites of humans, livestock, and crops, e.g the coffee borer beetle, hessian fly, head lice, and the citrus mealybug. Improved understanding of the evolutionary consequences of these unusual lifecycles and genetics therefore also has practical relevance in guiding our use of chemical, biological, and genetic controls.

## Supplementary Material

This is contained in a separate document.

## Acknowledgements

We thank J. Rayner, K. Stucky, S. Sturiale, L. Yusuf, and the Ross lab for helpful discussion.

