## Supplementary Material for "Sexual antagonism in haplodiploids"

### Outline

This Supplementary Material contains background on the models, and additional results, for those conditions presented in the main text of *Sexual antagonism in haplodiploids*. The document is organised as follows. **Section 1** summarises the general background and associated notation for the life-cycles and genetic systems analysed. **Section 2** provides details of the invasion analysis of a sexually antagonistic allele across these different life-cycles and genetic systems. **Section 3** is a kin-selection analysis of a sexually antagonistic trait using the methodology of Taylor and Frank (1996). Together these approaches provide complementary insights. The invasion analysis allows us to consider arbitrary strength of selection, and the effects of specific genetic parameters, such as dominance. The latter, by honing in on additivity and weak selection, both aids in interpretation of the invasion conditions and allows us to relax some of the assumptions about the ecology.

### 1 General Background

#### 1.1 Life cycles

We consider three core life-cycle variants that apply to most of our genetic systems: a fixed sib-mating scenario, a viscous population with mating occuring prior to dispersal, and a viscous population where mating occurs after dispersal. These three scenarios are described below and illustrated in Figure S1. In all three cases we describe a infinite population, split into a large number of patches (i.e. an infinite island model (Wright, 1931)). Across these cases, on each patch there is one, singly mated, female breeder, who lays a large brood of $\kappa$ offspring, of which of which a proportion $1-z$ are female, and $z$ are male. We also consider two further life-cycles which are specific to particular genetic systems: oedipal mating, and hermaphroditism in *Icerya purchasi*.

**Sib-mating scenario.** In the fixed sib-mating scenario, with probability $s$ females mate with brothers, and with probability $1-s$ mate with males from the population at large. This captures scenarios where there may be a temporal separation between sib-mating and outbreeding, analogous to both "prior selfing" and "delayed selfing" models in hermaphroditic plants, where there is no direct competition between pollen from self and from other plants (Lloyd and Schoen, 1992; Lloyd, 1992; Jordan and Connallon, 2014). After mating, females then either disperse with probability $d_f$, or remain on their natal patch with probability $1-d_f$. They then compete for the breeding spot on the patch, unsuccessful females then die, and the life-cycle begins again.

**Viscous population, mating before female dispersal.** In the mating pre-dispersal scenario females lay a brood of offspring as above, however now males first either disperse with probability $d_m$ or remain on their natal patch with probability $1-d_m$, females then mate with the males on their patch. Thus this is analogous to "mass-action" models of selfing in hermaphroditic plants where an individual's own pollen does directly compete with pollen from the wider population (Holsinger, 1991; Jordan and Connallon, 2014). After mating, females either disperse with probability $d_f$, or remain on their natal patch with probability $1-d_f$. Once again the females compete for the breeding spot on the patch, unsuccessful females then die, and the life-cycle begins anew.

**Viscous population, mating post female dispersal.** In the mating post-dispersal scenario, females once again lay a large brood of offspring as above. Now, however, both males and females disperse from the patch, with probabilities $d_m$ and $d_f$ respectively, or remain on the natal patch with probabilities $1-d_m$ and $1-d_f$ respectively. Females then mate with males on their patch, before competing for breeding spots. Unsuccessful females then die, and the life-cycle begins once more. This life-cycle is very similar to that previously analysed by Flintham and colleagues (Flintham, Savolainen, and Mullon, 2021).

**Oedipal mating.** For arrhenotokous organisms, females need not mate in order to produce offspring, and this opens up further variants of the life-cycles above. One particular case is oedipal mating, which has been documented in a range of arrhenotokous organisms (references in main text). In these cases a virgin female may initially lay a brood of exclusively male offspring, mating with one of her sons, before laying a bisexual brood. To incorporate this, we allow a proportion $\mathcal{O}$ of females to oedipally mate, and a proportion $1-\mathcal{O}$ to mate with a random male in the population. We assume that there is no selection that occurs amongst the first male brood, as is consistent with the ecology of many of these organisms, where a female simply mates with the first male who ecloses.

**Pseudo-hermaphroditism in Icerya.** *Icerya purchasi* is among the few "hermaphroditic" species of insect that are known, with diploid females and haploid males. However, in addition to these two genomes, in females there is a third genome belonging to an invasive lineage of spermatogenic tissue. This tissue can both fertilise the eggs of the female, but is also directly transmitted to female offspring. Females therefore can either "self" mating with this invasive male tissue, or "outbreed" mating with a true male. We allow for this by setting a proportion $\varphi$ of females to self, and proportion $1-\varphi$ mate with males as normal.

#### 1.2 Genetic systems

In the main text we described four main types of genetic system: diploidy, male paternal genome elimination (PGE), arrhenotoky, and pseudo-hermaphroditism in Icerya. Here we consider three further genetic systems: haploidy, whereby both males and females are haploid, female maternal genome elimination (MGE), where females are initially diploid but eliminate their maternal-origin genes when producing eggs, and paterothylotoky, where females are haploid producing either empty eggs which become fertilised to become females or haploid eggs which become fertilised to become males.

Of these wider sets of systems, only haploidy is well described and observed in nature, found in groups of green algae, brown algae, and bryophytes (Bachtrog et al., 2014; Coelho et al., 2018). Whilst examples of paterothylotoky have not been observed, these results are equivalent to those for the non-pseudoautosomal region of Z chromosomes in ZW chromosome systems. Finally, again whilst there are no known species that exhibit female MGE, there are various documented examples of androgenesis, whereby genes contributed by females are eliminated shortly after fertilisation (Burt and Trivers, 2006; Schwander and Oldroyd, 2016), thus a sex-specific version of this is not inconceivable. Moreover, these systems provide a comparison to investigate and distinguish between the effects of sex-specific aspects of ecology and genetics on the invasion conditions.

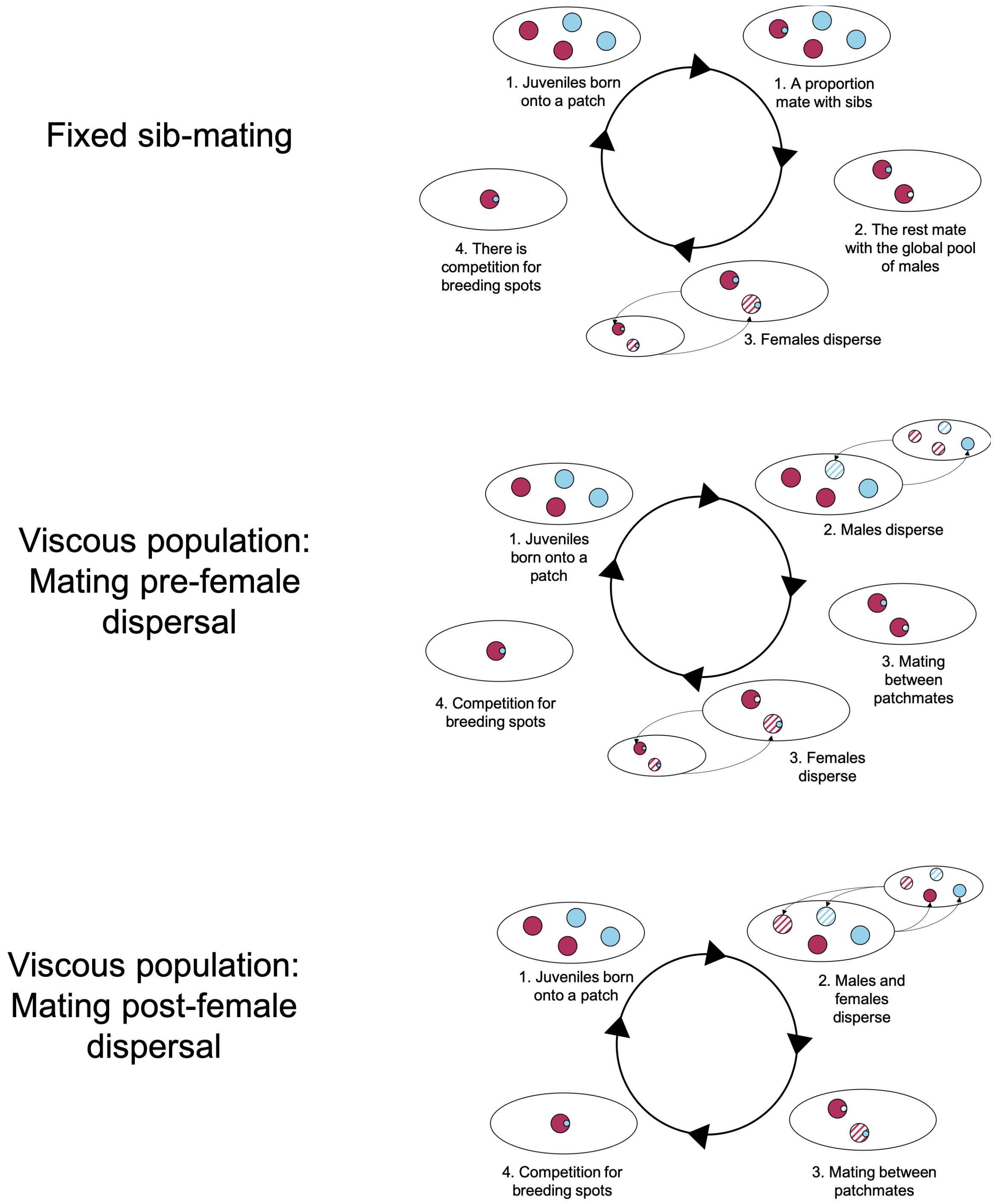

Figure S1: The structure of the three main life-cycles that we consider

### 2 Invasion analysis

For the invasion analysis we first write out recursion equations describing the frequency of different mating pairs. We then make weak selection approximations of these recursion equations, and use these to analytically find the invasion conditions for a sexually antagonistic allele. Finally, we use these recursion equations to find the invasion boundaries for stronger selection regimes, and thus the parameter space for stable polymorphisms.

#### 2.1 Notation

We consider the conditions when a mutant allele will be able to invade a population which is monomorphic for a resident allele. For a particular genetic system, we write out the genotype of a female as $a_i$ where $a_i \in A$, with $A$ being the set of all possible female genotypes in that system, and the genotype of a male as $b_i$, where $b_i \in B$, with $B$ being the set of all possible male genotypes, and the genotype of a mating pair $c_i$, with $c_i \in C$, where $C$ is the set of all possible mating pairs.

We write out the frequency of a particular mating pair genotype at time $t$ as $f_{c_i}$, and the frequency at time $t+1$ to be $f'_{c_i}$. The number of females of genotype $a_j$ produced by mating pair $c_i$ is given by $x_{a_j,c_i}$, and the number of males of genotype $k$ produced by genotype $c_i$ is $y_{b_k,c_i}$. Let $w_{a_j,c_i}$ be the fitness of an individual female who has genotype $a_j$ and who comes from mating pair of genotype $c_i$, and let $v_{b_k,c_i}$ be the fitness of an individual male with genotype $b_k$ and who comes from a mating pair with genotype $c_i$. We write out the mean fitness of the females produced by a mating pair of genotype $c_i$ to be $\overline{w}_{c_i} = (1/x_{c_i})\sum_j x_{a_j,c_i} w_{a_j,c_i}$, and the mean fitness of the males produced by a mating pair of genotype $c_i$ to be $\overline{v}_{c_i} = (1/y_{c_i})\sum_k y_{b_k,c_i} v_{b_k,c_i}$, where the number of females produced by a genotype $c_i$ is $x_{c_i} = \sum_j x_{a_j,c_i}$ and the number of males $y_{c_i} = \sum_k y_{b_k,c_i}$.

#### 2.2 Methodology

##### 2.2.1 Recursion equations

**Fixed sib-mating scenario**

$$\begin{aligned} f'_{\{a_j,b_k\}} = \sum_i f_{c_i} [(1-d_f) \left( \frac{x_{a_j,c_i} w_{a_j,c_i}}{\alpha_i} (s \frac{y_{b_k,c_i} v_{b_k,c_i}}{y_{c_i} \overline{v}_{c_i}} + (1-s) \sum_l f_{c_l} \frac{y_{b_k,c_l} v_{b_k,c_l}}{\overline{yv}}) \right) \\ + d_f \sum_{\mathbb{i}} f_{c_{\mathbb{i}}} \frac{x_{a_j,c_i} w_{a_j,c_i}}{\alpha_{\mathbb{i}}} (s \frac{y_{b_k,c_i} v_{b_k,c_i}}{y_{c_i} \overline{v}_{c_i}} + (1-s) \sum_l f_{c_l} \frac{y_{b_k,c_l} v_{b_k,c_l}}{\overline{yv}})] \end{aligned} \tag{S2.2.1a}$$

Where $\alpha_i$ is the competitiveness of patch type $c_i$ for females:

$$\alpha_i = (1-d_f) x_{c_i} \overline{w}_{c_i} + d_f \overline{xw} \tag{S2.2.1b}$$

And where the the mean male and female fitnesses are:

$$\overline{w} = \sum_i f_{c_i} \overline{w}_{c_i} = \sum_i f_{c_i} \sum_j \frac{x_{a_j,c_i}}{x_{c_i}} w_{a_j,c_i} \tag{S2.2.1c}$$

$$\overline{v} = \sum_i f_{c_i} \overline{v}_{c_i} = \sum_i f_{c_i} \sum_k \frac{y_{b_k,c_i}}{y_{c_i}} v_{b_k,c_i} \tag{S2.2.1d}$$

**Viscous population, mating pre-female dispersal**

$$
\begin{aligned}
f'_{\{a_j,b_k\}} = &\sum_i f_{c_i}[(1-d_f)\left(\frac{x_{a_j,c_i}w_{a_j,c_i}}{\alpha_i}((1-d_m)\frac{y_{b_k,c_i}v_{b_k,c_i}}{\beta_i}+d_m\sum_l f_{c_l}\frac{y_{b_k,c_l}v_{b_k,c_l}}{\beta_i})\right) \\
&+d_f\sum_{\mathbb{i}} f_{c_{\mathbb{i}}}\frac{x_{a_j,c_i}w_{a_j,c_i}}{\alpha_{\mathbb{i}}}((1-d_m)\frac{y_{b_k,c_i}v_{b_k,c_i}}{\beta_i}+d_m\sum_l f_{c_l}\frac{y_{b_k,c_l}v_{b_k,c_l}}{\beta_i})]
\end{aligned} \tag{S2.2.2a}
$$

$$\alpha_i = (1-d_f)x_{c_i}\overline{w}_{c_i} + d_f\overline{xw} \tag{S2.2.2b}$$

And the relative competitiveness of patch type $c_i$ for males $\beta_i$ is:

$$\beta_i = (1-d_m)y_{c_i}\overline{v}_{c_i} + d_m\overline{yv} \tag{S2.2.2c}$$

With the mean female and male fitnesses given by:

$$\overline{w} = \sum_i f_{c_i}\overline{w}_{c_i} = \sum_i f_{c_i}\sum_j \frac{x_{a_j,c_i}}{x_{c_i}}w_{a_j,c_i} \tag{S2.2.2d}$$

$$\overline{v} = \sum_i f_{c_i}\overline{v}_{c_i} = \sum_i f_{c_i}\sum_k \frac{y_{b_k,c_i}}{y_{c_i}}v_{b_k,c_i} \tag{S2.2.2e}$$

**Viscous population, mating post-female dispersal**

$$
\begin{aligned}
f'_{\{a_j,b_k\}} = &\sum_i f_{c_i}[(1-d_f)\left(\frac{x_{a_j,c_i}w_{a_j,c_i}}{\alpha_i}((1-d_m)\frac{y_{b_k,c_i}v_{b_k,c_i}}{\beta_i}+d_m\sum_l f_{c_l}\frac{y_{b_k,c_l}v_{b_k,c_l}}{\beta_i})\right) \\
&+d_f\sum_{\mathbb{i}} f_{c_{\mathbb{i}}}\frac{x_{a_j,c_i}w_{a_j,c_i}}{\alpha_{\mathbb{i}}}((1-d_m)\frac{y_{b_k,c_{\mathbb{i}}}v_{b_k,c_{\mathbb{i}}}}{\beta_{\mathbb{i}}}+d_m\sum_l f_{c_l}\frac{y_{b_k,c_l}v_{b_k,c_l}}{\beta_{\mathbb{i}}})]
\end{aligned} \tag{S2.2.3a}
$$

With the competitiveness of patch type $c_i$ for males and females given by:

$$\alpha_i = (1-d_f)x_{c_i}\overline{w}_{c_i} + d_f\overline{xw} \tag{S2.2.3b}$$

$$\beta_i = (1-d_m)y_{c_i}\overline{v}_{c_i} + d_m\overline{yv} \tag{S2.2.3c}$$

With the mean female and male fitnesses given by:

$$\overline{w} = \sum_i f_{c_i}\overline{w}_{c_i} = \sum_i f_{c_i}\sum_j \frac{x_{a_j,c_i}}{x_{c_i}}w_{a_j,c_i} \tag{S2.2.3d}$$

$$\overline{v} = \sum_i f_{c_i}\overline{v}_{c_i} = \sum_i f_{c_i}\sum_k \frac{y_{b_k,c_i}}{y_{c_i}}v_{b_k,c_i} \tag{S2.2.3e}$$

**Oedipal mating** We notate the fraction of oedipal mating to be $\mathcal{O}$. Also, as when a female mates she produces a brood of offspring herself, we write out the number of these males of genotype $b_k$ to be $y_{b_k,a_j}$, and her total number of males to be $y_{a_j}$.

$$
\begin{aligned}
f'_{\{a_j,b_k\}} = &\sum_i f_{c_i}[(1-d_f)\left(\frac{x_{a_j,c_i}w_{a_j,c_i}}{\alpha_i}(\mathcal{O}\frac{y_{b_k,a_j}}{y_{a_j}}+(1-\mathcal{O})\sum_l f_{c_l}\frac{y_{b_k,c_l}v_{b_k,c_l}}{\overline{yv}})\right) \\
&+d_f\sum_{\mathbb{i}} f_{c_{\mathbb{i}}}\frac{x_{a_j,c_i}w_{a_j,c_i}}{\alpha_{\mathbb{i}}}(\mathcal{O}\frac{y_{b_k,a_j}}{y_{a_j}}+(1-\mathcal{O})\sum_l f_{c_l}\frac{y_{b_k,c_l}v_{b_k,c_l}}{\overline{yv}})]
\end{aligned} \tag{S2.2.4a}
$$

Where $\alpha_i$ is the competitiveness of patch type $c_i$ for females:

$$\alpha_i = (1-d_f)x_{c_i}\overline{w}_{c_i} + d_f\overline{xw} \tag{S2.2.4b}$$

And where the the mean male and female fitnesses are:

$$\overline{w} = \sum_i f_{c_i}\overline{w}_{c_i} = \sum_i f_{c_i}\sum_j \frac{x_{a_j,c_i}}{x_{c_i}} w_{a_j,c_i} \tag{S2.2.4c}$$

$$\overline{v} = \sum_i f_{c_i}\overline{v}_{c_i} = \sum_i f_{c_i}\sum_k \frac{y_{b_k,c_i}}{y_{c_i}} v_{b_k,c_i} \tag{S2.2.4d}$$

**Pseudo-hermaphroditism in Icerya** We notate the fraction of females who are mated by the infectious haploid spermatogenic tissue to to be $\varphi$, and the proportion mated by 'true' males to be $1-\varphi$. Similar to in the oedipal mating scenario, when a female 'selfs' we imagine that she produces a brood of males of which she then mates with one, in this case we notate it $\mathbb{y}_{b_k,a_j}$, as is distinct from the genotypes of 'true' males she would produce.

$$\begin{aligned} f'_{\{a_j,b_k\}} = \sum_i f_{c_i}[(1-d_f)\left(\frac{x_{a_j,c_i}w_{a_j,c_i}}{\alpha_i}(\varphi\frac{\mathbb{y}_{b_k,a_j}}{\mathbb{y}_{a_j}} + (1-\varphi)\sum_l f_{c_l}\frac{y_{b_k,c_l}v_{b_k,c_l}}{\overline{yv}})\right) \\ + d_f\sum_{\mathbb{i}} f_{c_{\mathbb{i}}}\frac{x_{a_j,c_i}w_{a_j,c_i}}{\alpha_{\mathbb{i}}}(\varphi\frac{\mathbb{y}_{b_k,a_j}}{\mathbb{y}_{a_j}} + (1-\varphi)\sum_l f_{c_l}\frac{y_{b_k,c_l}v_{b_k,c_l}}{\overline{yv}})] \end{aligned} \tag{S2.2.5a}$$

Where $\alpha_i$ is the competitiveness of patch type $c_i$ for females:

$$\alpha_i = (1-d_f)x_{c_i}\overline{w}_{c_i} + d_f\overline{xw} \tag{S2.2.5b}$$

And where the the mean male and female fitnesses are:

$$\overline{w} = \sum_i f_{c_i}\overline{w}_{c_i} = \sum_i f_{c_i}\sum_j \frac{x_{a_j,c_i}}{x_{c_i}} w_{a_j,c_i} \tag{S2.2.5c}$$

$$\overline{v} = \sum_i f_{c_i}\overline{v}_{c_i} = \sum_i f_{c_i}\sum_k \frac{y_{b_k,c_i}}{y_{c_i}} v_{b_k,c_i} \tag{S2.2.5d}$$

##### 2.2.2 Jacobians

Using these recursion equations we can ask when the mutant allele will be able to invade from rarity. If we denote the mating pair purely made up the resident allele $c_*$, then we want to consider the equilibrium point $f_{c_*} = 1$. If this is unstable then the mutant allele mating pair genotypes will be able to invade. To determine the stability, we first calculate the Jacobian matrix $\mathbf{J}$, analysed when the mating pair genotypes containing the mutant allele are vanishingly rare in the population (Otto and Day, 2011). Each entry of the matrix is given by:

$$\mathbf{J}_{a,b} = \frac{\partial f'_{c_a}}{\partial f'_{c_b}}\Big|_{c_*=1} \tag{S2.2.6}$$

If the leading eigenvalue of this matrix is greater than one, $\lambda_{max} > 1$ then the mutant mating pairs will increased in frequency, and thus the mutant allele will be able to invade.

##### 2.2.3 Weak selection approximations

For many of these scenarios it is not tractable to find full analytical solutions. Instead we approximate the conditions for invasion when the mutant allele has a vanishingly small fitness effect as compared to the resident allele. If this is the case our fitness effects are of order $\delta$, and we can write the largest eigenvalue as:

$$\lambda_{max} \approx 1 + \delta\lambda \tag{S2.2.7}$$

We then substitute this back into our characteristic equation and perform a first order Taylor expansion, ignoring terms of order $\delta^2$ and higher. We then solve for $\lambda$, and compute our condition for invasion. The invasion conditions for the sib-mating, DMD, and DDM life-cycles can be found in Tables S2-S4. The invasion conditions for the oedipdal mating and the pseudo-hermaphroditism of Icerya can be found in Table S5.

##### 2.2.4 Numerical solutions

We also find numerical approximations for the invasion boundary under stronger selection regimes by first finding the largest eigenvalue for specific values, and then interpolating the boundary where $\lambda_{max} = 1$ by using Mathematica's ListContour function (Inc., n.d.). These results can be seen in Figures S5-S7.

#### 2.3 Invasion conditions

| | Rice: Female beneficial | Rice: Male beneficial | Kidwell |
|---|---|---|---|
| $w_0$ | 1 | 1 | $1-u_f$ |
| $w_1$ | $1+S$ | $1-T$ | 1 |
| $w_{00}$ | 1 | 1 | $1-u_f$ |
| $w_{01}$ | $1+h_fS$ | $1-Th_f$ | $1-h_fu_f$ |
| $w_{10}$ | $1+h_fS$ | $1-Th_f$ | $1-h_fu_f$ |
| $w_{11}$ | $1+S$ | $1-T$ | 1 |
| $v_0$ | 1 | 1 | 1 |
| $v_1$ | $1-T$ | $1+S$ | $1-u_m$ |
| $v_{00}$ | 1 | 1 | 1 |
| $v_{01}$ | $1-h_mT$ | $1+h_mS$ | $1-h_mu_m$ |
| $v_{10}$ | $1-h_mT$ | $1+h_mS$ | $1-h_mu_m$ |
| $v_{11}$ | $1-T$ | $1+S$ | $1-u_m$ |

Table S1: Fitness scheme used to analyse sexual antagonism. In the two Rice scenarios, $S$ represents the benefit of the allele, $T$ the cost, and $h_f$ and $h_m$ are the dominance coefficients in males and females respectively (Rice 1984). In the Kidwell scenario, the fittest genotype has fitness 1, with the cost $u_f$ to females and $u_m$ to males. Again $h_f$ and $h_m$ are the dominance coefficients in males and females.

| | Fixed Inbreeding | Viscous Population: Mating pre-dispersal | Viscous Population: Mating post-dispersal |
|---|---|---|---|
| Haploidy | $\frac{T}{S} < \frac{(s+1)\phi_f^2-2}{2(s-1)}$ | $\frac{T}{S} < \frac{\phi_f^2(\phi_m+1)-2}{\phi_m^2+\phi_m-2}$ | $\frac{T}{S} < \frac{\phi_f(\phi_f+\phi_m)-2}{\phi_m(\phi_f+\phi_m)-2}$ |
| Diploidy | $\frac{T}{S} < \frac{(4(s-1)h_f-s)\left((s+1)\phi_f^2-2\right)}{8(s-1)^2h_m-2(s-1)s}$ | $\frac{T}{S} < \frac{\left(\phi_f^2(\phi_m+1)-2\right)(4h_f(\phi_m-1)-\phi_m)}{(\phi_m-1)(\phi_m+2)(4h_m(\phi_m-1)-\phi_m)}$ | $\frac{T}{S} < \frac{(\phi_f(\phi_f+\phi_m)-2)((4h_f-1)\phi_f\phi_m-4h_f)}{(\phi_m(\phi_f+\phi_m)-2)(\phi_f(4h_m-1)\phi_m-4h_m)}$ |
| Arrhenotoky | $\frac{T}{S} < -\frac{(4(s-1)h_f-s)\left(3\phi_f^2+s-4\right)}{(s-4)(s-2)(s-1)}$ | $\frac{T}{S} < -\frac{\left(3\phi_f^2+\phi_m-4\right)(4h_f(\phi_m-1)-\phi_m)}{(\phi_m-2)(\phi_m-1)((\phi_m-1)\phi_m-4)}$ | $\frac{T}{S} < \frac{\left(\left(\phi_f^2-2\right)\phi_f\phi_m-3\phi_f^2+4\right)((4h_f-1)\phi_f\phi_m-4h_f)}{(\phi_f\phi_m-2)(\phi_f(\phi_m^2-3)\phi_m-2\phi_m^2+4)}$ |
| Paterothylotoky | $\frac{T}{S} < -\frac{(s-2)\left((s+2)\phi_f^2+s-4\right)}{(s-4)(s-1)(4(s-1)h_m-s)}$ | $\frac{T}{S} < -\frac{(\phi_m-2)\left(\phi_f^2(\phi_m+2)+\phi_m-4\right)}{(\phi_m-1)((\phi_m-2)\phi_m-4)(4h_m(\phi_m-1)-\phi_m)}$ | $\frac{T}{S} < \frac{(\phi_f\phi_m-2)\left(\left(\phi_f^2-3\right)\phi_f\phi_m-2\phi_f^2+4\right)}{(\phi_f(\phi_m^2-2)\phi_m-3\phi_m^2+4)(\phi_f(4h_m-1)\phi_m-4h_m)}$ |
| Male PGE | $\frac{T}{S} < \frac{(4(s-1)h_f-s)\left(3\phi_f^2+s-4\right)}{8(s-1)^2h_m-2(s-1)s}$ | $\frac{T}{S} < \frac{\left(3\phi_f^2+\phi_m-4\right)(4h_f(\phi_m-1)-\phi_m)}{(\phi_m-1)(\phi_m+2)(4h_m(\phi_m-1)-\phi_m)}$ | $\frac{T}{S} < -\frac{\left(\left(\phi_f^2-2\right)\phi_f\phi_m-3\phi_f^2+4\right)((4h_f-1)\phi_f\phi_m-4h_f)}{(\phi_m(\phi_f+\phi_m)-2)(\phi_f(4h_m-1)\phi_m-4h_m)}$ |
| Female PGE | $\frac{T}{S} < \frac{(4(s-1)h_f-s)\left(((s-3)s-1)\phi_f^2-(s-2)s+2\right)}{(s-4)(s-1)(4(s-1)h_m-s)}$ | $\frac{T}{S} < \frac{\left(\phi_f^2((\phi_m-3)\phi_m-1)-(\phi_m-2)\phi_m+2\right)(4h_f(\phi_m-1)-\phi_m)}{(\phi_m-1)((\phi_m-2)\phi_m-4)(4h_m(\phi_m-1)-\phi_m)}$ | $\frac{T}{S} < -\frac{(\phi_f(\phi_f+\phi_m)-2)((4h_f-1)\phi_f\phi_m-4h_f)}{(\phi_f(\phi_m^2-2)\phi_m-3\phi_m^2+4)(\phi_f(4h_m-1)\phi_m-4h_m)}$ |

Table S2: Invasion conditions for female beneficial alleles under various genetic systems, and in various life-cycle structures, when the trait is under offspring control, and selection is weak

| | Fixed Inbreeding | Viscous Population: Mating pre-dispersal | Viscous Population: Mating post-dispersal |
|---|---|---|---|
| Haploidy | $\frac{T}{S} < \frac{(s+1)\left(\phi_f^2-1\right)}{s-1}$ | $\frac{T}{S} < \frac{\phi_f^2-1}{\phi_m-1}$ | $\frac{T}{S} < \frac{\phi_f^2-1}{\phi_m^2-1}$ |
| Diploidy | $\frac{T}{S} < \frac{(s+1)\left(\phi_f^2-1\right)(4(s-1)h_f-s)}{(s-1)(4(s-1)h_m-s)}$ | $\frac{T}{S} < \frac{\left(\phi_f^2-1\right)(4h_f(\phi_m-1)-\phi_m)}{(\phi_m-1)(4h_m(\phi_m-1)-\phi_m)}$ | $\frac{T}{S} < \frac{\left(\phi_f^2-1\right)((4h_f-1)\phi_f\phi_m-4h_f)}{(\phi_m^2-1)(\phi_f(4h_m-1)\phi_m-4h_m)}$ |
| Arrhenotoky | $\frac{T}{S} < \frac{((s-2)s-2)\left(\phi_f^2-1\right)(4(s-1)h_f-s)}{(s-2)(s-1)(4(s-1)h_m-s)}$ | $\frac{T}{S} < \frac{\left(\phi_f^2-1\right)((\phi_m-2)\phi_m-2)(4h_f(\phi_m-1)-\phi_m)}{(\phi_m-2)(\phi_m-1)(\phi_m+1)(4h_m(\phi_m-1)-\phi_m)}$ | $\frac{T}{S} < -\frac{2\left(\phi_f^2-1\right)((4h_f-1)\phi_f\phi_m-4h_f)}{(\phi_m^2-1)(\phi_f\phi_m-2)(\phi_f(4h_m-1)\phi_m-4h_m)}$ |
| Paterothylotoky | $\frac{T}{S} < \frac{s(2s-5)\left(\phi_f^2-1\right)}{2(s-2)(s-1)}$ | $\frac{T}{S} < \frac{\left(\phi_f^2-1\right)\phi_m(2\phi_m-5)}{2(\phi_m-2)(\phi_m-1)(\phi_m+1)}$ | $\frac{T}{S} < -\frac{\phi_f\left(\phi_f^2-1\right)\phi_m}{2(\phi_m^2-1)(\phi_f\phi_m-2)}$ |
| Male PGE | $\frac{T}{S} < \frac{((s-2)s-2)\left(\phi_f^2-1\right)(4(s-1)h_f-s)}{(s-2)(s-1)(4(s-1)h_m-s)}$ | $\frac{T}{S} < \frac{\left(\phi_f^2-1\right)((\phi_m-2)\phi_m-2)(4h_f(\phi_m-1)-\phi_m)}{(\phi_m-2)(\phi_m-1)(\phi_m+1)(4h_m(\phi_m-1)-\phi_m)}$ | $\frac{T}{S} < -\frac{2\left(\phi_f^2-1\right)((4h_f-1)\phi_f\phi_m-4h_f)}{(\phi_m^2-1)(\phi_f\phi_m-2)(\phi_f(4h_m-1)\phi_m-4h_m)}$ |
| Female PGE | $\frac{T}{S} < \frac{s(s^2-7)\left(\phi_f^2-1\right)(4(s-1)h_f-s)}{(s-1)((s-1)s-4)(4(s-1)h_m-s)}$ | $\frac{T}{S} < \frac{\left(\phi_f^2-1\right)\phi_m(\phi_m^2-7)(4h_f(\phi_m-1)-\phi_m)}{(\phi_m-1)(\phi_m+1)((\phi_m-1)\phi_m-4)(4h_m(\phi_m-1)-\phi_m)}$ | $\frac{T}{S} < \frac{\phi_f\left(\phi_f^2-1\right)\phi_m(\phi_f\phi_m-3)((4h_f-1)\phi_f\phi_m-4h_f)}{(\phi_m^2-1)(\phi_f\phi_m(\phi_f\phi_m-1)-4)(\phi_f(4h_m-1)\phi_m-4h_m)}$ |

Table S3: Invasion conditions for female beneficial alleles under various genetic systems, and in various life-cycle structures, when the trait is under maternal control, and selection is weak

| | Fixed Inbreeding | Viscous Population: Mating pre-dispersal | Viscous Population: Mating post-dispersal |
|---|---|---|---|
| Haploidy | $\frac{T}{S} < \frac{(s+1)\left(\phi_f^2-1\right)}{s-1}$ | $\frac{T}{S} < \frac{\phi_f^2-1}{\phi_m-1}$ | $\frac{T}{S} < \frac{\phi_f^2-1}{\phi_m^2-1}$ |
| Diploidy | $\frac{T}{S} < \frac{(s+1)\left(\phi_f^2-1\right)(4(s-1)h_f-s)}{(s-1)(4(s-1)h_m-s)}$ | $\frac{T}{S} < \frac{\left(\phi_f^2-1\right)(4h_f(\phi_m-1)-\phi_m)}{(\phi_m-1)(4h_m(\phi_m-1)-\phi_m)}$ | $\frac{T}{S} < \frac{\left(\phi_f^2-1\right)((4h_f-1)\phi_f\phi_m-4h_f)}{(\phi_m^2-1)(\phi_f(4h_m-1)\phi_m-4h_m)}$ |
| Arrhenotoky | $\frac{T}{S} < \frac{((s-2)s+4)\left(\phi_f^2-1\right)}{(s-1)s}$ | $\frac{T}{S} < \frac{\left(\phi_f^2-1\right)((\phi_m-2)\phi_m+4)}{\phi_m(\phi_m^2-1)}$ | $\frac{T}{S} < -\frac{2\left(\phi_f^2-1\right)(\phi_f\phi_m-2)}{\phi_f\phi_m(\phi_m^2-1)}$ |
| Paterothylotoky | $\frac{T}{S} < \frac{(s+2)\left(\phi_f^2-1\right)(4(s-1)h_f-s)}{8(s-1)^2h_m-2(s-1)s}$ | $\frac{T}{S} < \frac{\left(\phi_f^2-1\right)(\phi_m+2)(4h_f(\phi_m-1)-\phi_m)}{2(\phi_m^2-1)(4h_m(\phi_m-1)-\phi_m)}$ | $\frac{T}{S} < -\frac{\left(\phi_f^2-1\right)(\phi_f\phi_m-2)((4h_f-1)\phi_f\phi_m-4h_f)}{2(\phi_m^2-1)(\phi_f(4h_m-1)\phi_m-4h_m)}$ |
| Male PGE | $\frac{T}{S} < \frac{(s((s-2)s-1)-4)\left(\phi_f^2-1\right)(4(s-1)h_f-s)}{(s-3)(s-1)s(4(s-1)h_m-s)}$ | $\frac{T}{S} < \frac{\left(\phi_f^2-1\right)(\phi_m((\phi_m-2)\phi_m-1)-4)(4h_f(\phi_m-1)-\phi_m)}{(\phi_m-3)(\phi_m-1)\phi_m(\phi_m+1)(4h_m(\phi_m-1)-\phi_m)}$ | $\frac{T}{S} < \frac{\left(\phi_f^2-1\right)(\phi_f\phi_m(\phi_f\phi_m-1)-4)((4h_f-1)\phi_f\phi_m-4h_f)}{\phi_f\phi_m(\phi_m^2-1)(\phi_f\phi_m-3)(\phi_f(4h_m-1)\phi_m-4h_m)}$ |
| Female PGE | $\frac{T}{S} < \frac{(s+2)\left(\phi_f^2-1\right)(4(s-1)h_f-s)}{8(s-1)^2h_m-2(s-1)s}$ | $\frac{T}{S} < \frac{\left(\phi_f^2-1\right)(\phi_m+2)(4h_f(\phi_m-1)-\phi_m)}{2(\phi_m^2-1)(4h_m(\phi_m-1)-\phi_m)}$ | $\frac{T}{S} < -\frac{\left(\phi_f^2-1\right)(\phi_f\phi_m-2)((4h_f-1)\phi_f\phi_m-4h_f)}{2(\phi_m^2-1)(\phi_f(4h_m-1)\phi_m-4h_m)}$ |

Table S4: Invasion conditions for female beneficial alleles under various genetic systems, and in various life-cycle structures, when the trait is under paternal control, and selection is weak

| | Invasion conditions |
|---|---|
| Oedipal mating | $\frac{\tau}{\sigma} < -\frac{(2(\upsilon-1)h_f-\upsilon)((\upsilon+3)\phi_f^2-4)}{4(\upsilon-1)}$ |
| Pseudo-hermaphroditism in Icerya | $\frac{\tau}{\sigma} < \frac{(4(s-1)(\varphi-1)h_f+s(-\varphi)+s+2\varphi)(\phi_f^2(s(\varphi-1)\varphi+\varphi^2-3)+s((3-2\varphi)\varphi-1)-2\varphi+4)}{(s-1)(\varphi-1)(s(\varphi-1)+2)(s(\varphi-1)-2\varphi+4)}$ |

Table S5: Invasion conditions for female beneficial alleles under various genetic systems, and in various life-cycle structures, when the trait is under offspring control, and selection is weak

#### 2.4 Potential for polymorphism

| | Conditions for stable polymorphism |
|---|---|
| Haploidy | n/a |
| Diploidy | $-\frac{2(s-1)(4(s-1)h_m-3s+4)}{(4(s-1)h_f-s)((s+1)\phi_f^2-2)} < \frac{u_f}{u_m} < \frac{2(s-1)(s-4(s-1)h_m)}{(4(s-1)h_f-3s+4)((s+1)\phi_f^2-2)}$ |
| Arrhenotoky | $-\frac{(s-4)(s-2)(s-1)}{(4(s-1)h_f-s)(3\phi_f^2+s-4)} < \frac{u_f}{u_m} < \frac{(s-4)(s-2)(s-1)}{(4(s-1)h_f-3s+4)(3\phi_f^2+s-4)}$ |
| Paterothylotoky | $\frac{(s-4)(s-1)(4(s-1)h_m-3s+4)}{(s-2)((s+2)\phi_f^2+s-4)} < \frac{u_f}{u_m} < \frac{(s-4)(s-1)(s-4(s-1)h_m)}{(s^2-4)\phi_f^2+s^2-6s+8}$ |
| Male PGE | $-\frac{2(s-1)(4(s-1)h_m-3s+4)}{(4(s-1)h_f-s)(3\phi_f^2+s-4)} < \frac{u_f}{u_m} < \frac{2(s-1)(s-4(s-1)h_m)}{(4(s-1)h_f-3s+4)(3\phi_f^2+s-4)}$ |
| Female MGE | $-\frac{(s-4)(s-1)(4(s-1)h_m-3s+4)}{(4(s-1)h_f-s)(((s-3)s-1)\phi_f^2-(s-2)s+2)} < \frac{u_f}{u_m} < \frac{(s-4)(s-1)(s-4(s-1)h_m)}{(4(s-1)h_f-3s+4)(((s-3)s-1)\phi_f^2-(s-2)s+2)}$ |

Table S6: Weak selection approximations for conditions for the maintenance of a stable polymorphism with sib-mating under various genetic systems, when the trait is under offspring control

| | Conditions for stable polymorphism |
|---|---|
| Haploidy | n/a |
| Diploidy | $-\frac{(\phi_m-1)(\phi_m+2)(4h_m(\phi_m-1)-3\phi_m+4)}{(\phi_f^2(\phi_m+1)-2)(4h_f(\phi_m-1)-\phi_m)} < \frac{u_f}{u_m} < -\frac{(\phi_m-1)(\phi_m+2)(4h_m(\phi_m-1)-\phi_m)}{(\phi_f^2(\phi_m+1)-2)(4h_f(\phi_m-1)-3\phi_m+4)}$ |
| Arrhenotoky | $-\frac{(\phi_m-2)(\phi_m-1)((\phi_m-1)\phi_m-4)}{(3\phi_f^2+\phi_m-4)(4h_f(\phi_m-1)-\phi_m)} < \frac{u_f}{u_m} < \frac{(\phi_m-2)(\phi_m-1)((\phi_m-1)\phi_m-4)}{(3\phi_f^2+\phi_m-4)(4h_f(\phi_m-1)-3\phi_m+4)}$ |
| Paterothylotoky | $\frac{(\phi_m-1)((\phi_m-2)\phi_m-4)(4h_m(\phi_m-1)-3\phi_m+4)}{(\phi_m-2)(\phi_f^2(\phi_m+2)+\phi_m-4)} < \frac{u_f}{u_m} < -\frac{(\phi_m-1)((\phi_m-2)\phi_m-4)(4h_m(\phi_m-1)-\phi_m)}{(\phi_m-2)(\phi_f^2(\phi_m+2)+\phi_m-4)}$ |
| Male PGE | $-\frac{(\phi_m-1)(\phi_m+2)(4h_m(\phi_m-1)-3\phi_m+4)}{(3\phi_f^2+\phi_m-4)(4h_f(\phi_m-1)-\phi_m)} < \frac{u_f}{u_m} < -\frac{(\phi_m-1)(\phi_m+2)(4h_m(\phi_m-1)-\phi_m)}{(3\phi_f^2+\phi_m-4)(4h_f(\phi_m-1)-3\phi_m+4)}$ |
| Female MGE | $-\frac{(\phi_m-1)((\phi_m-2)\phi_m-4)(4h_m(\phi_m-1)-3\phi_m+4)}{(\phi_f^2((\phi_m-3)\phi_m-1)-(\phi_m-2)\phi_m+2)(4h_f(\phi_m-1)-\phi_m)} < \frac{u_f}{u_m} < -\frac{(\phi_m-1)((\phi_m-2)\phi_m-4)(4h_m(\phi_m-1)-\phi_m)}{(\phi_f^2((\phi_m-3)\phi_m-1)-(\phi_m-2)\phi_m+2)(4h_f(\phi_m-1)-3\phi_m+4)}$ |

Table S7: Weak selection approximations for the conditions for the maintenance of a stable polymorphism with mating pre-female dispersal under various genetic systems, when the trait is under offspring control

| | Conditions for stable polymorphism |
|---|---|
| Haploidy | n/a |
| Diploidy | $-\frac{(\phi_m(\phi_f+\phi_m)-2)(\phi_f(4h_m-3)\phi_m-4h_m+4)}{(\phi_f(\phi_f+\phi_m)-2)((4h_f-1)\phi_f\phi_m-4h_f)} < \frac{u_f}{u_m} < -\frac{(\phi_m(\phi_f+\phi_m)-2)(\phi_f(4h_m-1)\phi_m-4h_m)}{(\phi_f(\phi_f+\phi_m)-2)((4h_f-3)\phi_f\phi_m-4h_f+4)}$ |
| Arrhenotoky | $\frac{(\phi_f\phi_m-2)(\phi_f(\phi_m^2-3)\phi_m-2\phi_m^2+4)}{\left(\left(\phi_f^2-2\right)\phi_f\phi_m-3\phi_f^2+4\right)((4h_f-1)\phi_f\phi_m-4h_f)} < \frac{u_f}{u_m} < -\frac{(\phi_f\phi_m-2)(\phi_f(\phi_m^2-3)\phi_m-2\phi_m^2+4)}{\left(\left(\phi_f^2-2\right)\phi_f\phi_m-3\phi_f^2+4\right)((4h_f-3)\phi_f\phi_m-4h_f+4)}$ |
| Paterothylotoky | $-\frac{(\phi_f(\phi_m^2-2)\phi_m-3\phi_m^2+4)(\phi_f(4h_m-3)\phi_m-4h_m+4)}{(\phi_f\phi_m-2)\left(\left(\phi_f^2-3\right)\phi_f\phi_m-2\phi_f^2+4\right)} < \frac{u_f}{u_m} < \frac{(\phi_f(\phi_m^2-2)\phi_m-3\phi_m^2+4)(\phi_f(4h_m-1)\phi_m-4h_m)}{(\phi_f\phi_m-2)\left(\left(\phi_f^2-3\right)\phi_f\phi_m-2\phi_f^2+4\right)}$ |
| Male PGE | $\frac{(\phi_m(\phi_f+\phi_m)-2)(\phi_f(4h_m-3)\phi_m-4h_m+4)}{\left(\left(\phi_f^2-2\right)\phi_f\phi_m-3\phi_f^2+4\right)((4h_f-1)\phi_f\phi_m-4h_f)} < \frac{u_f}{u_m} < \frac{(\phi_m(\phi_f+\phi_m)-2)(\phi_f(4h_m-1)\phi_m-4h_m)}{\left(\left(\phi_f^2-2\right)\phi_f\phi_m-3\phi_f^2+4\right)((4h_f-3)\phi_f\phi_m-4h_f+4)}$ |
| Female MGE | $\frac{(\phi_f(\phi_m^2-2)\phi_m-3\phi_m^2+4)(\phi_f(4h_m-3)\phi_m-4h_m+4)}{(\phi_f(\phi_f+\phi_m)-2)((4h_f-1)\phi_f\phi_m-4h_f)} < \frac{u_f}{u_m} < \frac{(\phi_f(\phi_m^2-2)\phi_m-3\phi_m^2+4)(\phi_f(4h_m-1)\phi_m-4h_m)}{(\phi_f(\phi_f+\phi_m)-2)((4h_f-3)\phi_f\phi_m-4h_f+4)}$ |

Table S8: Weak selection approximations for conditions for the maintenance of a stable polymorphism with mating post-female dispersal under various genetic systems, when the trait is under offspring control

#### 2.5 Figures

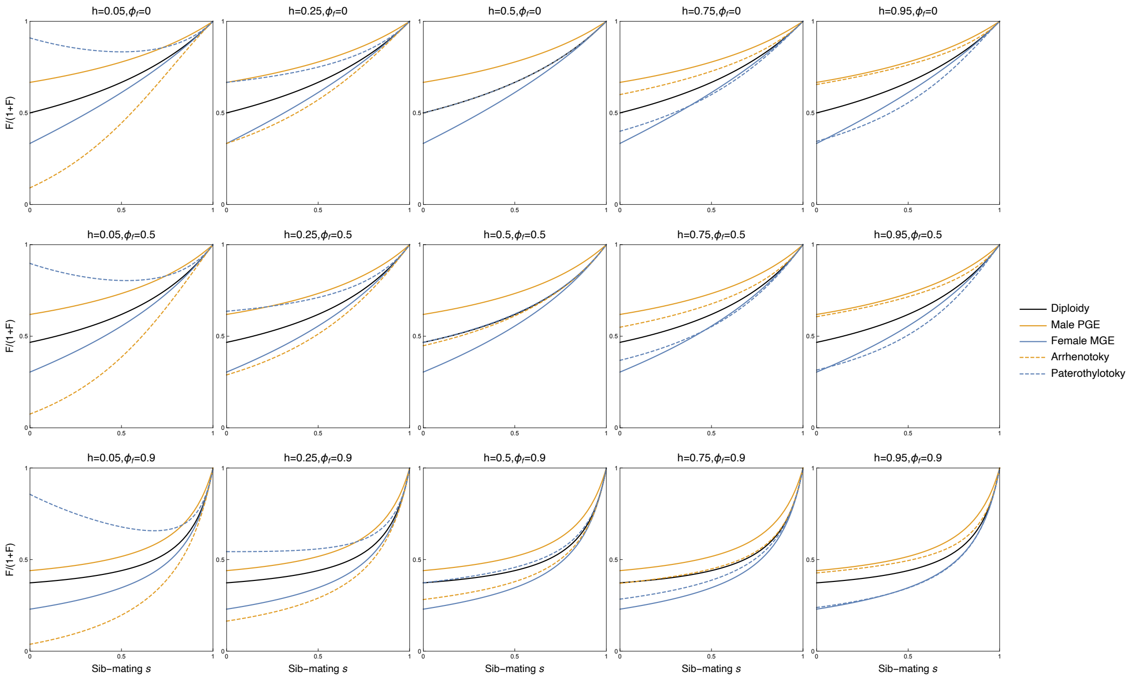

Figure S2: Potential for feminisation across some of the different genetic systems we consider as a function of dominance $h$, sib-mating $s$, and female philopatry $\phi_f$

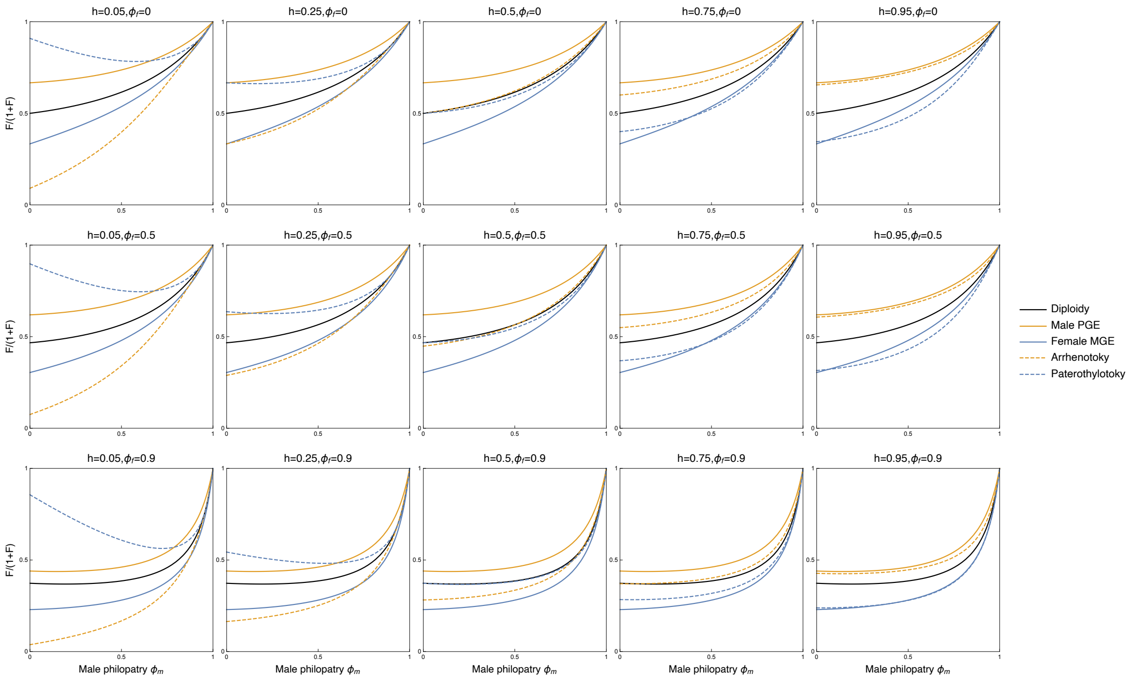

Figure S3: Potential for feminisation across some of the different genetic systems we consider when mating occurs pre-female dispersal, as a function of dominance $h$, male philopatry $\phi_m$, and female philopatry $\phi_f$

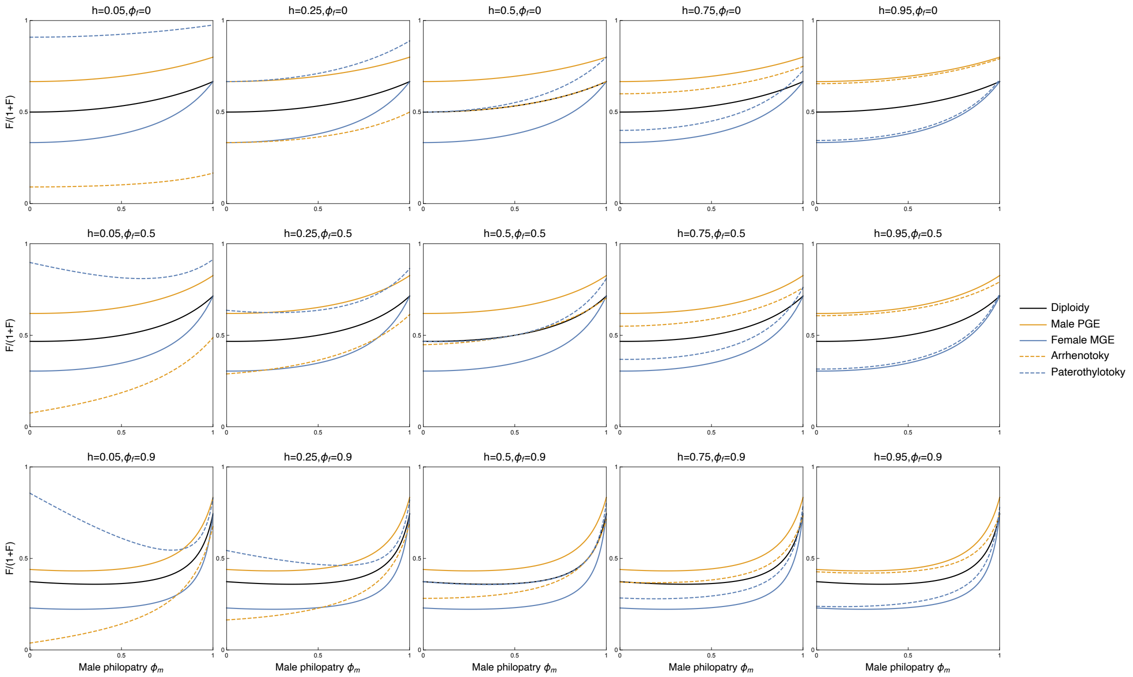

Figure S4: Potential for feminisation across some of the different genetic systems we consider when mating occurs post-female dispersal, as a function of dominance $h$, male philopatry $\phi_m$, and female philopatry $\phi_f$

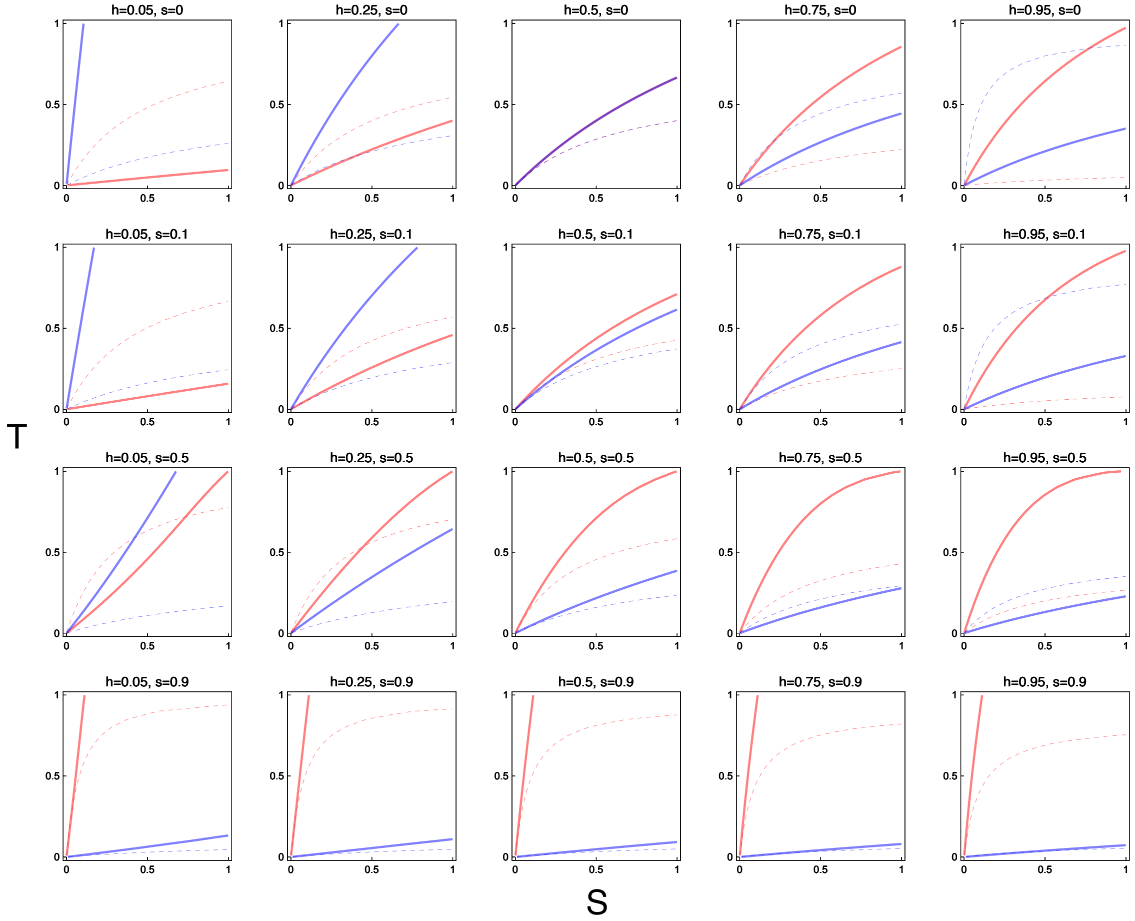

Figure S5: Potential for polymorphism for male and female beneficial alleles under arrhenotoky with sib-mating, across different levels of sib-mating $s$, and dominance coefficients $h$, where dominance is parallel across sexes $h = h_f = h_m$

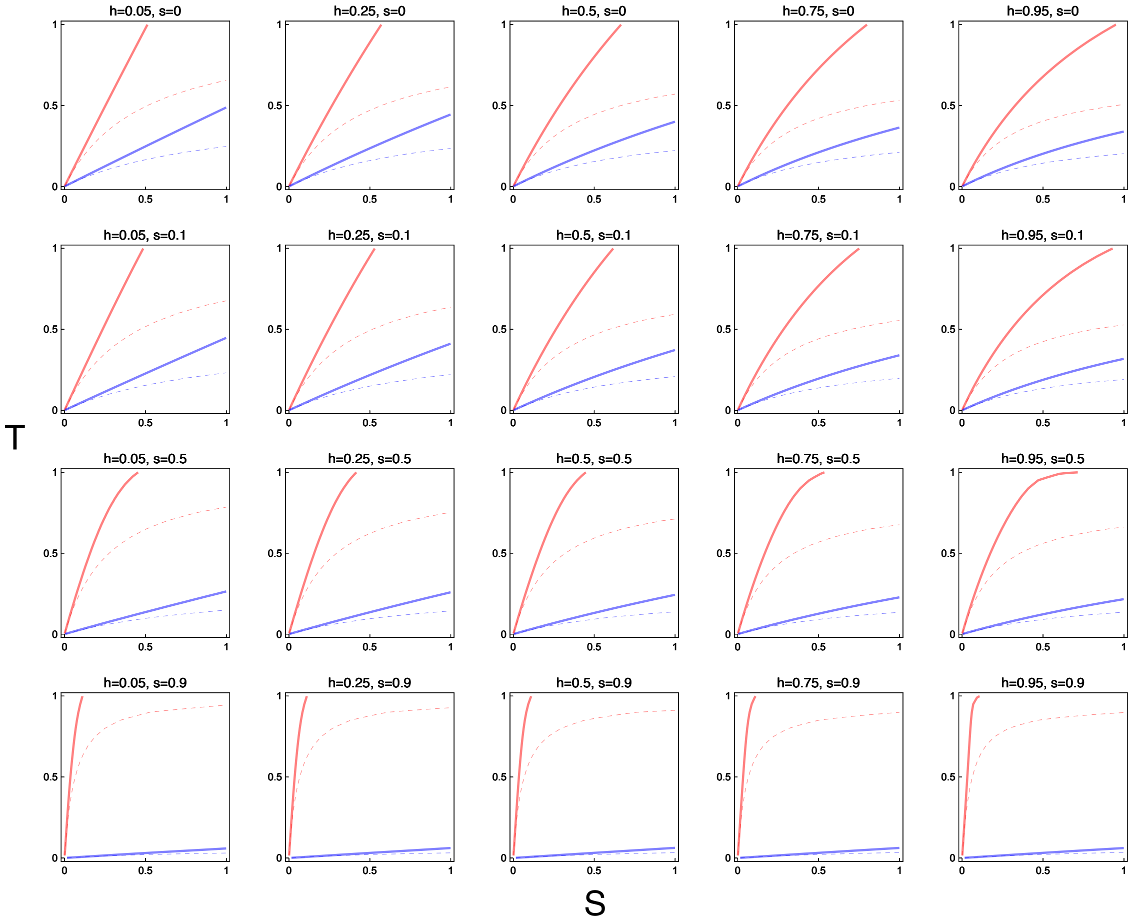

Figure S6: Potential for polymorphism for male and female beneficial alleles under male PGE with sib-mating, across different levels of sib-mating $s$, and dominance coefficients $h$, where dominance is parallel across sexes $h = h_f = h_m$

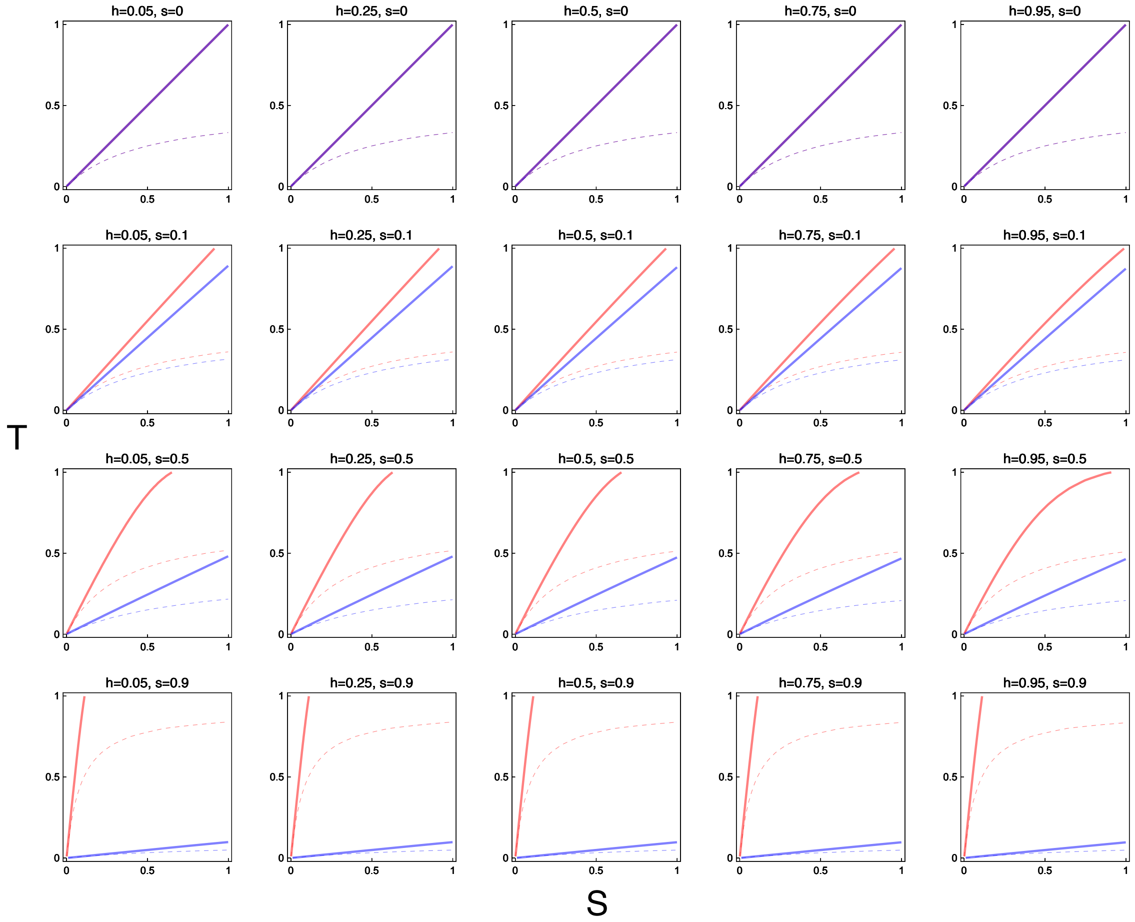

Figure S7: Potential for polymorphism for male and female beneficial alleles under diploidy with sib-mating, across different levels of sib-mating $s$, and dominance coefficients $h$, where dominance is parallel across sexes $h = h_f = h_m$

### 3 Kin selection model of a sexually antagonistic trait

Alongside the above invasion analysis, we also perform a kin-selection analysis of a sexually antagonistic trait $\phi$, using the direct fitness methodology of Taylor and Frank (1996). By assuming additivity and weak selection, we can more straightforwardly partition the different effects of the above life-cycles on sexual antagonism into their component parts. Doing this enables a more general analysis of how different life-cycle structures and mating schemes may impact sexual antagonism. We also then use this to understand how, assuming specific fitness functions, this will alter the optimum of such traits, and how, depending on who controls such traits, the optima reached may differ.

#### 3.1 General conditions

We represent the relative fitness of an individual in our population by $W$, where $W$ is function of $\boldsymbol{\phi}$, a vector of the different values of our sexually antagonistic trait $\phi$ in our population, including our focal individual. We consider a locus which affects this trait, and denote the genic value of a gene drawn at random from the population at this locus $g$, and the population average of this trait value to be $\overline{g}$. The condition for natural selection to favour an increase in the level of this sexually antagonistic trait is:

$$\frac{dW}{dg}\bigg|_{g=\overline{g}} > 0 \tag{S3.1.1a}$$

And similarly for a decrease in this trait value:

$$\frac{dW}{dg}\bigg|_{g=\overline{g}} < 0 \tag{S3.1.1b}$$

If our population is composed of different types of individual, as it is the cases we investigate, then these individuals may be differently affected by changes in the trait value. We write the relative fitness of a focal individual of class $j$ as $W_j$. Furthermore, these different types of individual may, on average, make differential contributions to the ancestry of the population. To account for this, we must weight the effects in these different classes of individual by that class's reproductive value $c_j$, which is the product of the relative abundance of that class $u_j$, and the expected contribution of individuals of that class to the future ancestry of the population $v_j$, hence $c_j = u_j v_j$. These class reproductive values are normalised such that $\sum_j c_j = 1$. Doing this, our condition for increase becomes:

$$\frac{dW}{dg} = \sum_j c_j \frac{dW_j}{dg_j} > 0 \tag{S3.1.2}$$

Where $g_j$ denotes the genic value of a gene picked randomly at the locus in a focal individual of class $j$, and once again the condition is evaluated when the population is monomorphic for the trait value $g_j = g = \overline{g}$. Using the chain rule, we can expand the $dW_j/dg_j$ terms, separating the direct effects of the an individual's trait value upon their own fitness, and the effects of the trait values of other social partners upon their fitness. If we index individuals in our population by $i$ then:

$$\frac{dW_j}{dg_j} = \sum_i \frac{\partial W_j}{\partial \phi_i} \frac{d\phi_i}{dG_i} \frac{dG_i}{dg_i} \frac{dg_i}{dg_j} \tag{S3.1.3}$$

Where $d\phi_i/dG_i = \gamma$ is the mapping of breeding value to phenotype, which we assume to be a constant across our individuals, $dG_i/dg_i$ is the change in an individual's genetic value with a change in the genetic value at a single locus, which is 1 under adding genetics and $1/n_i$ under averaging genetics with $n_i$ the ploidy of the $i$th individual (Frank, 2003; Gardner, 2012), and $dg_i/dg_j = \rho_{i,j}$ is the consanguinity between our focal individual of class $j$, and the individual $i$. Our total condition for increase can then be written for averaging genetics as:

$$\frac{dW}{dg} = \sum_j \sum_i c_j \frac{\partial W_j}{\partial \phi_i} \frac{\rho_{i,j}}{n_i} > 0 \tag{S3.1.4a}$$

And for adding genetics:

$$\frac{dW}{dg} = \sum_j \sum_i c_j \frac{\partial W_j}{\partial \phi_i} \rho_{i,j} > 0 \tag{S3.1.4b}$$

**Further class structure** The above analysis assumes that the different genes within an individual contribute equally to the genic value of the individual that they reside within, and also are transmitted equally from that individual. However, this may not be always be the case, for example if there is genomic imprinting, then one of the two parent-of-origin copies may solely determine the phenotype, or alternatively if there is paternal-genome elimination, then only one of the maternal-origin gene copy will be transmitted. To account for this, we now further decompose our population into different classes of genes, where $j$ now indexes a focal gene in class $j$, for example "female, maternal-origin", and $i$ indexes all of the individual gene copies in our population, $\mathbb{i}$ indexes the individuals in the population, and $W$ is now the relative fitness of an individual gene copy, rather than individual. Our condition for increase now becomes:

$$\frac{dW_j}{dg_j} = \sum_i \frac{\partial W_j}{\partial \phi_{\mathbb{i}}} \frac{d\phi_{\mathbb{i}}}{dG_{\mathbb{i}}} \frac{dG_{\mathbb{i}}}{dg_i} \frac{dg_i}{dg_j} \tag{S3.1.5}$$

Much remains the same as before, however now the term $dG_{\mathbb{i}}/dg_i$ captures the relative effect of a specific gene copy $i$ on it's individual's genetic value $G_{\mathbb{i}}$, rather than being an average across the gene copies in that individual. For instance, if there is genomic imprinting, with the paternal-origin copy silenced, then if that gene copy is of maternal-origin then $dG_{\mathbb{f}}/dg_{f^{mat}} = 1$, whilst if that gene is of paternal-origin $dG_{\mathbb{f}}/dg_{f^{pat}} = 0$. Similarly, a male paternal-origin gene's fitness will be altered differently to that of a male maternal-origin gene under paternal-genome elimination, i.e. $\partial W_{m^{pat}}/\partial \phi_{\mathbb{i}} = 0$.

#### 3.2 Fitness functions

**General model** We can describe the above three life-cycles above with the following model, where $W$ is the relative fitness of an individual female, and $V$ the relative fitness of an individual male. The competitiveness of a female $w$, and a male $v$, are functions of a sexually antagonistic trait $\phi$. We denote the phenotype of our focal female as $\phi_x^f$, our focal male as $\phi_x^m$, female and male patchmates as $\phi_y^f$ and $\phi_y^m$, and of females and males in the wider population as $\phi_z^f$ and $\phi_z^m$.

Individuals may compete with both the global pool of individuals, or with individuals to whom they are related. We can capture the variation between these extremes with the term $a$, the spatial scale of competition (Frank, 1998). $a_{ff}$ is the scale of competition for females upon related females, $a_{mm}$ is the scale of competition for males upon related males, and $a_{fm}$ is the scale of competition for females upon related males. Individuals may also mate with individuals from the global pool or with relatives, the probability that an individual mates with a patchmate is given by $\psi$, and the probability that they mate with an individual from the rest of the population is $(1-\psi)$. The values for the scale of competition that emerge from the different life-cycles described can be seen in Table S9.

$$W, W_{Mat}, W_{Pat} = \frac{w[\phi_x^f]}{a_{ff}\, w[\phi_y^f] + (1-a_{ff})\, w[\phi_z^f]} \tag{S3.2.1a}$$

$$V, V_{Mat}, V_{Pat} = \left(\frac{v[\phi_x^m]}{a_{mm}\, v[\phi_y^m] + (1-a_{mm})\, v[\phi_z^m]}\right)\left(\frac{\psi\, w[\phi_y^f] + (1-\psi)\, w[\phi_z^f]}{a_{fm}\, w[\phi_y^f] + (1-a_{fm})\, w[\phi_z^f]}\right) \tag{S3.2.1b}$$

**Specific Gaussian model** The above model leaves open the specific relationship between the sexually antagonistic trait and competitiveness. We construct a toy model where the competitiveness of females $w$ and males $v$ are two Gaussian functions of the trait $\phi$, normally distributed around the female and male optima, $\phi^{f*}$ and $\phi^{m*}$ respectively, with standard deviations $SD_f$ and $SD_m$ respectively:

$$w[\phi] = N[\phi^{f*}, SD_f] \tag{S3.2.2a}$$

$$v[\phi] = N[\phi^{m*}, SD_m] \tag{S3.2.2b}$$

#### 3.3 Genotype to phenotype mapping

We now look at how the genetic values at different gene positions of an individual $g$ map into the breeding value of the individual $G$, and therefore the trait value $\phi$ of that individual. For haploid systems there is only a single gene determining the phenotype, and therefore it is straightforwardly determined by this gene copy. For diploid systems (including arrhenotoky and paterothylotoky), multiple gene positions may contribute to the phenotype, and these different gene positions may have different amounts of influence on the breeding value. We assign a fraction $\mathscr{X}$ of the breeding value in females to the maternal-origin gene, and fraction $(1-\mathscr{X})$ to the paternal-origin gene. In males, we assign a fraction $(1-\mathscr{Y})$ of the breeding value to the maternal-origin gene and a fraction $\mathscr{Y}$ to the paternal-origin gene. Earlier we also discussed how the choice of adding vs averaging genetics is important when comparing across ploidy levels. To allow for this choice we weight the breeding value in females and males by $\alpha$ and $\beta$ respectively. In all cases there is an uncorrelated error denoted by $\epsilon$.

Additionally, it may be that the trait value $\phi$ of our focal individual is not necessarily under the control of that individual themselves, but instead is determined by the breeding value of another individual. In particular here, we consider the influence of parents. We denote the value of the parents of our focal individual by $\mathbb{x}$. Thus the maternal-origin gene of our focal individual's mother would be denoted $\mathbb{x}_{f_{Mat}}$, and the paternal-origin gene of this focal individual's father by $\mathbb{x}_{m_{Pat}}$.

##### 3.3.1 Haploidy

**Offspring control**

$$\phi_x^f = G_x^f + \epsilon = g_{x,f} + \epsilon \tag{S3.3.1a}$$

$$\phi_x^m = G_x^m + \epsilon = g_{x,m} + \epsilon \tag{S3.3.1b}$$

**Maternal control**

$$\phi_x^f = G_{\mathbb{X}}^f + \epsilon = g_{\mathbb{X},f} + \epsilon \tag{S3.3.2a}$$

$$\phi_x^m = G_{\mathbb{X}}^f + \epsilon = g_{\mathbb{X},f} + \epsilon \tag{S3.3.2b}$$

**Paternal control**

$$\phi_x^f = G_{\mathbb{X}}^m + \epsilon = g_{\mathbb{X},m} + \epsilon \tag{S3.3.3a}$$

$$\phi_x^m = G_{\mathbb{X}}^m + \epsilon = g_{\mathbb{X},m} + \epsilon \tag{S3.3.3b}$$

##### 3.3.2 Diploidy

**Offspring control**

$$\phi_x^f = G_x^f + \epsilon = \alpha\left(\mathscr{X} g_{x,f_{Mat}} + (1-\mathscr{X}) g_{x,f_{Pat}}\right) + \epsilon \tag{S3.3.4a}$$

$$\phi_x^m = G_x^m + \epsilon = \beta\left((1-\mathscr{Y}) g_{x,m_{Mat}} + \mathscr{Y} g_{x,m_{Pat}}\right) + \epsilon \tag{S3.3.4b}$$

**Maternal control**

$$\phi_x^f = G_{\mathbb{X}}^f + \epsilon = \alpha\left(\mathscr{X} g_{\mathbb{X},f_{Mat}} + (1-\mathscr{X}) g_{\mathbb{X},f_{Pat}}\right) + \epsilon \tag{S3.3.5a}$$

$$\phi_x^m = G_{\mathbb{X}}^f + \epsilon = \alpha\left(\mathscr{X} g_{\mathbb{X},f_{Mat}} + (1-\mathscr{X}) g_{\mathbb{X},f_{Pat}}\right) + \epsilon \tag{S3.3.5b}$$

**Paternal control**

$$\phi_x^m = G_{\mathbb{X}}^m + \epsilon = \beta\left((1-\mathscr{Y}) g_{\mathbb{X},m_{Mat}} + \mathscr{Y} g_{\mathbb{X},m_{Pat}}\right) + \epsilon \tag{S3.3.6a}$$

$$\phi_x^m = G_{\mathbb{X}}^m + \epsilon = \beta\left((1-\mathscr{Y}) g_{\mathbb{X},m_{Mat}} + \mathscr{Y} g_{\mathbb{X},m_{Pat}}\right) + \epsilon \tag{S3.3.6b}$$

| | $a_{ff}$ | $a_{fm}$ | $a_{mm}$ |
|---|---|---|---|
| Fixed sib-mating | $(1-d_f)^2$ | $(1-d_f)^2$ | $s$ |
| Mating pre-dispersal | $(1-d_f)^2$ | $(1-d_f)^2$ | $(1-d_m)^2$ |
| Mating post-dispersal | $(1-d_f)^2$ | $(1-d_m)(1-d_f)$ | $(1-d_m)^2$ |

Table S9: The sex-specific scales of competition in our three different life-cycles

| | $\partial W_{\text{ff}}$ | $\partial W_{\text{fm}}$ | $\partial W_{\text{mf}}$ | $\partial W_{\text{mm}}$ |
|---|---|---|---|---|
| $\partial x_{\text{ff}}$ | $\alpha\sigma\mathscr{X}$ | $\alpha\sigma\mathscr{X}$ | 0 | 0 |
| $\partial y_{\text{ff}}$ | $\alpha\sigma(-\mathscr{X})a_{\text{ff}}$ | $\alpha\sigma(-\mathscr{X})a_{\text{ff}}$ | $\alpha\sigma\mathscr{X}(\psi - a_{\text{fm}})$ | $\alpha\sigma\mathscr{X}(\psi - a_{\text{fm}})$ |
| $\partial x_{\text{fm}}$ | $\alpha\sigma(1-\mathscr{X})$ | $\alpha\sigma(1-\mathscr{X})$ | 0 | 0 |
| $\partial y_{\text{fm}}$ | $\alpha\sigma(-(1-\mathscr{X}))a_{\text{ff}}$ | $\alpha\sigma(-(1-\mathscr{X}))a_{\text{ff}}$ | $\alpha\sigma(1-\mathscr{X})(\psi - a_{\text{fm}})$ | $\alpha\sigma(1-\mathscr{X})(\psi - a_{\text{fm}})$ |
| $\partial x_{\text{mf}}$ | 0 | 0 | $\beta\tau(1-\mathscr{Y})$ | $\beta\tau(1-\mathscr{Y})$ |
| $\partial y_{\text{mf}}$ | 0 | 0 | $\beta\tau(-(1-\mathscr{Y}))a_{\text{mm}}$ | $\beta\tau(-(1-\mathscr{Y}))a_{\text{mm}}$ |
| $\partial x_{\text{mm}}$ | 0 | 0 | $\beta\tau\mathscr{Y}$ | $\beta\tau\mathscr{Y}$ |
| $\partial y_{\text{fm}}$ | 0 | 0 | $\beta\tau(-\mathscr{Y})a_{\text{mm}}$ | $\beta\tau(-\mathscr{Y})a_{\text{mm}}$ |

Table S10: Marginal fitness effects for different genetic actors on self and social partners under diploidy.

#### 3.4 Marginal fitness effects

With these fitness functions, and the mapping of genotype to phenotype, we can now look at how the fitness of our focal individual changes with the changing genetic values of self and social partners. To get the marginal fitness effect of different actors, we can differentiate our fitness functions by the values of the various gene positions, evaluated when there is vanishingly small genetic variation, $g_{x,i} = g_{y,i} = g_{z,i} = \overline{g}$.

For the general model, we notate the marginal change in the competitiveness in females with respect to the trait value by:

$$\frac{dw[\phi]/d\phi}{w[\phi]} = \sigma \tag{S3.4.1a}$$

And in males:

$$\frac{dv[\phi]/d\phi}{v[\phi]} = \tau \tag{S3.4.1b}$$

When a trait is sexually antagonistic, $\tau$ and $\sigma$ will have opposite signs. The values of $\sigma$ and $\tau$ will depend on the specific mapping of our trait into competitiveness. In our toy model:

$$\sigma = \left(\phi^{f*} - \alpha\phi^{f}\right)\frac{1}{SD_f^2} \tag{S3.4.2a}$$

$$\tau = \left(\phi^{m*} - \beta\phi^{m}\right)\frac{1}{SD_m^2} \tag{S3.4.2b}$$

The marginal fitness effects under haploidy, diploidy, and under offspring, maternal, and paternal control, can be seen in Table S10.

#### 3.5 Consanguinities

To calculate the consanguinities between different gene positions, we first write out recursions describing the probability of identity by descent (Bulmer, 1994). We then assume that the consanguinity coefficients have reached their quasi-equilibrium values which is a reasonable assumption if selection is weak (Gardner, West, and Wild, 2011).

We denote the probability of being IBD between two gene positions within an individual as $\rho^{i}_{g_1,g_2}$, and the probability of being IBD between two gene positions within a patch as $\rho^{p}_{g_1,g_2}$. We denote the probability of being IBD between a mother and an offspring $\rho^{Mot}_{g_1,g_2}$, where $g_1$ is the gene position in the mother, and $g_2$ the gene position in the offspring. We denote the probability of being IBD between a father and offspring $\rho^{Fat}_{g_1,g_2}$, where $g_1$ is the gene position in the father, and $g_2$ the gene position in the offspring.

The probability that a maternal-origin gene came from a maternal-origin gene is $\mathscr{A}$, and the probability that a paternal-origin gene came from a paternal-origin gene is $\mathscr{B}$. Such that for diploidy $\mathscr{A} = 1/2, \mathscr{B} = 1/2$, for arrhenotoky/male PGE $\mathscr{A} = 1/2, \mathscr{B} = 0$, and for paterothylotoky/female PGE $\mathscr{A} = 0, \mathscr{B} = 1/2$. The probability that an individual descended from a mating between patchmates is $\psi$. The values for $\psi$ for our different life-cycles/mating systems are: $\phi = s$ for sib-mating, $\phi = (1 - d_m)$ for DMD, and $\phi = (1 - d_m)(1 - d_f)$ for DDM.

The consanguinities between these gene positions under these different genetic systems can be found in Table S11 . (ADD in parental/haploid tables).

##### 3.5.1 Haploidy

**Within individuals**

$$\rho^{i}_{f} = \rho^{i}_{m} = 1 \tag{S3.5.1}$$

**Between patchmates**

$$\rho^{p}_{f,f} = (1-\mathscr{A})(\mathscr{A}\rho^{p}_{f,m}\psi + (1-\mathscr{A})) + \mathscr{A}((1-\mathscr{A})\rho^{p}_{f,m}\psi + \mathscr{A}) \tag{S3.5.2a}$$

$$\rho^{p}_{f,m} = (1-\mathscr{A})(\rho^{p}_{f,m}\psi(1-\mathscr{B}) + \mathscr{B}) + \mathscr{A}(\rho^{p}_{f,m}\psi\mathscr{B} + (1-\mathscr{B})) \tag{S3.5.2b}$$

$$\rho^{p}_{m,m} = \mathscr{B}(\rho^{p}_{f,m}\psi(1-\mathscr{B}) + \mathscr{B}) + (1-\mathscr{B})(\rho^{p}_{f,m}\psi\mathscr{B} + (1-\mathscr{B})) \tag{S3.5.2c}$$

**Mothers and offspring**

$$\rho^{Mot}_{f,f} = (1-\mathscr{A})\rho^{p}_{f,m}\psi + \mathscr{A} \tag{S3.5.3a}$$

$$\rho^{Mot}_{f,m} = \rho^{p}_{f,m}\psi\mathscr{B} + (1-\mathscr{B}) \tag{S3.5.3b}$$

**Fathers and offspring**

$$\rho^{Fat}_{m,f} = \mathscr{A}\rho^{p}_{f,m}\psi + (1-\mathscr{A}) \tag{S3.5.4a}$$

$$\rho^{Fat}_{m,m} = \rho^{p}_{f,m}\psi(1-\mathscr{B}) + \mathscr{B} \tag{S3.5.4b}$$

##### 3.5.2 Diploidy

**Within individuals**

$$\rho^i_{ff,ff} = \rho^i_{fm,fm} = \rho^i_{mf,mf} = \rho^i_{mm,mm} = 1 \tag{S3.5.5a}$$

$$\rho^i_{ff,fm} = \psi\left(\mathscr{A}\left((1-\mathscr{B})\rho^p_{ff,mf} + \mathscr{B}\rho^p_{ff,mm}\right) + (1-\mathscr{A})\left((1-\mathscr{B})\rho^p_{fm,mf} + \mathscr{B}\rho^p_{fm,mm}\right)\right) \tag{S3.5.5b}$$

$$\rho^i_{mf,mm} = \psi\left(\mathscr{A}\left((1-\mathscr{B})\rho^p_{ff,mf} + \mathscr{B}\rho^p_{ff,mm}\right) + (1-\mathscr{A})\left((1-\mathscr{B})\rho^p_{fm,mf} + \mathscr{B}\rho^p_{fm,mm}\right)\right) \tag{S3.5.5c}$$

**Between patchmates**

$$\rho^p_{ff,ff} = \mathscr{A}\left(\mathscr{A}\rho^i_{ff,ff} + (1-\mathscr{A})\,\rho^i_{ff,fm}\right) + (1-\mathscr{A})\left(\mathscr{A}\rho^i_{ff,fm} + (1-\mathscr{A})\,\rho^i_{ff,ff}\right) \tag{S3.5.6a}$$

$$\rho^p_{ff,fm} = \psi\left(\mathscr{A}\left((1-\mathscr{B})\,\rho^p_{ff,mf} + \mathscr{B}\rho^p_{ff,mm}\right) + (1-\mathscr{A})\left((1-\mathscr{B})\,\rho^p_{fm,mf} + \mathscr{B}\rho^p_{fm,mm}\right)\right) \tag{S3.5.6b}$$

$$\rho^p_{ff,mf} = (1-\mathscr{A})\left(\rho^i_{ff,fm}\mathscr{A} + 1\,(1-\mathscr{A})\right) + \mathscr{A}\left(\rho^i_{ff,fm}\,(1-\mathscr{A}) + 1\mathscr{A}\right) \tag{S3.5.6c}$$

$$\rho^p_{ff,mm} = \psi\left(\mathscr{A}\left(\rho^p_{ff,mf}\,(1-\mathscr{B}) + \rho^p_{ff,mm}\mathscr{B}\right) + (1-\mathscr{A})\left(\rho^p_{fm,mf}\,(1-\mathscr{B}) + \rho^p_{fm,mm}\mathscr{B}\right)\right) \tag{S3.5.6d}$$

$$\rho^p_{fm,fm} = (1-\mathscr{B})\left(\rho^i_{mf,mm}\mathscr{B} + 1\,(1-\mathscr{B})\right) + \mathscr{B}\left(\rho^i_{mf,mm}\,(1-\mathscr{B}) + 1\mathscr{B}\right) \tag{S3.5.6e}$$

$$\rho^p_{fm,mf} = \psi\left((1-\mathscr{B})\left(\rho^p_{ff,mf}\mathscr{A} + \rho^p_{fm,mf}\,(1-\mathscr{A})\right) + \mathscr{B}\left(\rho^p_{ff,mm}\mathscr{A} + \rho^p_{fm,mm}\,(1-\mathscr{A})\right)\right) \tag{S3.5.6f}$$

$$\rho^p_{fm,mm} = (1-\mathscr{B})\left(\rho^i_{mf,mm}\mathscr{B} + 1\,(1-\mathscr{B})\right) + \mathscr{B}\left(\rho^i_{mf,mm}\,(1-\mathscr{B}) + 1\mathscr{B}\right) \tag{S3.5.6g}$$

$$\rho^p_{mf,mf} = (1-\mathscr{A})\left(\rho^i_{ff,fm}\mathscr{A} + 1\,(1-\mathscr{A})\right) + \mathscr{A}\left(\rho^i_{ff,fm}\,(1-\mathscr{A}) + 1\mathscr{A}\right) \tag{S3.5.6h}$$

$$\rho^p_{mf,mm} = \psi\left(\mathscr{A}\left(\rho^p_{ff,mf}\,(1-\mathscr{B}) + \rho^p_{ff,mm}\mathscr{B}\right) + (1-\mathscr{A})\left(\rho^p_{fm,mf}\,(1-\mathscr{B}) + \rho^p_{fm,mm}\mathscr{B}\right)\right) \tag{S3.5.6i}$$

$$\rho^p_{mm,mm} = (1-\mathscr{B})\left(\rho^i_{mf,mm}\mathscr{B} + 1\,(1-\mathscr{B})\right) + \mathscr{B}\left(\rho^i_{mf,mm}\,(1-\mathscr{B}) + 1\mathscr{B}\right) \tag{S3.5.6j}$$

**Mothers and offspring**

$$\rho^{Mat}_{ff,ff} = \rho^i_{ff,fm}\,(1-\mathscr{A}) + 1\mathscr{A} \tag{S3.5.7a}$$

$$\rho^{Mat}_{ff,fm} = \psi\left(\rho^p_{ff,mf}\,(1-\mathscr{B}) + \rho^p_{ff,mm}\mathscr{B}\right) \tag{S3.5.7b}$$

$$\rho^{Mot}_{ff,mf} = \rho^i_{ff,fm}\,(1-\mathscr{A}) + 1\mathscr{A} \tag{S3.5.7c}$$

$$\rho^{Mot}_{ff,mm} = \psi\left(\rho^{p}_{ff,mm}\mathscr{B} + \rho^{p}_{ff,mf}(1-\mathscr{B})\right) \tag{S3.5.7d}$$

$$\rho^{Mot}_{fm,ff} = \rho^{i}_{ff,fm}\mathscr{A} + 1\,(1-\mathscr{A}) \tag{S3.5.7e}$$

$$\rho^{Mot}_{fm,fm} = \psi\left(\rho^{p}_{fm,mf}(1-\mathscr{B}) + \rho^{p}_{fm,mm}\mathscr{B}\right) \tag{S3.5.7f}$$

$$\rho^{Mot}_{fm,mf} = \rho^{i}_{ff,fm}\mathscr{A} + 1\,(1-\mathscr{A}) \tag{S3.5.7g}$$

$$\rho^{Mot}_{fm,mm} = \psi\left(\rho^{p}_{fm,mf}(1-\mathscr{B}) + \rho^{p}_{fm,mm}\mathscr{B}\right) \tag{S3.5.7h}$$

**Fathers and offspring**

$$\rho^{Fat}_{mf,ff} = \psi\left(\rho^{p}_{ff,mf}\mathscr{A} + \rho^{p}_{fm,mf}(1-\mathscr{A})\right) \tag{S3.5.8a}$$

$$\rho^{Fat}_{mf,fm} = \rho^{i}_{mf,mm}\mathscr{B} + 1\,(1-\mathscr{B}) \tag{S3.5.8b}$$

$$\rho^{Fat}_{mf,mf} = \psi\left(\rho^{p}_{ff,mf}\mathscr{A} + \rho^{p}_{fm,mf}(1-\mathscr{A})\right) \tag{S3.5.8c}$$

$$\rho^{Fat}_{mf,mm} = \rho^{i}_{mf,mm}\mathscr{B} + 1\,(1-\mathscr{B}) \tag{S3.5.8d}$$

$$\rho^{Fat}_{mm,ff} = \psi\left(\rho^{p}_{ff,mm}\mathscr{A} + \rho^{p}_{fm,mm}(1-\mathscr{A})\right) \tag{S3.5.8e}$$

$$\rho^{Fat}_{mm,fm} = \rho^{i}_{mf,mm}(1-\mathscr{B}) + 1\mathscr{B} \tag{S3.5.8f}$$

$$\rho^{Fat}_{mm,mf} = \psi\left(\rho^{p}_{ff,mm}\mathscr{A} + \rho^{p}_{fm,mm}(1-\mathscr{A})\right) \tag{S3.5.8g}$$

$$\rho^{Fat}_{mm,mm} = \rho^{i}_{mf,mm}(1-\mathscr{B}) + 1\mathscr{B} \tag{S3.5.8h}$$

#### 3.6 Reproductive value

Reproductive value provides a measure of an individual gene's, or a class of genes', expected asymptotic contribution to future generations. We denote the reproductive value of class $i$ by $c_i$. This can be calculated by writing a gene flow matrix for a monomorphic population. This gene flow matrix is essentially describing a Markov process of a gene's state, going back in time (Grafen, 2006). We notate the probability that a gene in class $i$ came from class $j$ in the previous timestep by $\pi_{i,j}$. If we write this gene flow matrix out, then the dominant left eigenvector of this matrix gives us the class reproductive value weightings for our different gene

| | Haploidy |
|---|---|
| $\rho^{p}_{f,f}$ | $\frac{2\mathbb{L}^2-2\mathbb{L}+1}{2\mathbb{L}^2\psi-2\mathbb{L}\psi+1}$ |
| $\rho^{p}_{f,m}$ | $\frac{2\mathbb{L}^2-2\mathbb{L}+1}{2\mathbb{L}^2\psi-2\mathbb{L}\psi+1}$ |
| $\rho^{p}_{m,m}$ | $\frac{2\mathbb{L}^2-2\mathbb{L}+1}{2\mathbb{L}^2\psi-2\mathbb{L}\psi+1}$ |
| $\rho^{Mot}_{f,f}$ | $\frac{2\mathbb{L}^2\psi-\mathbb{L}\psi-\mathbb{L}+1}{2\mathbb{L}^2\psi-2\mathbb{L}\psi+1}$ |
| $\rho^{Mot}_{f,m}$ | $\frac{2\mathbb{L}^2\psi-\mathbb{L}\psi-\mathbb{L}+1}{2\mathbb{L}^2\psi-2\mathbb{L}\psi+1}$ |
| $\rho^{Fat}_{m,f}$ | $\frac{2\mathbb{L}^2\psi-3\mathbb{L}\psi+\mathbb{L}+\psi}{2\mathbb{L}^2\psi-2\mathbb{L}\psi+1}$ |
| $\rho^{Fat}_{m,m}$ | $\frac{2\mathbb{L}^2\psi-3\mathbb{L}\psi+\mathbb{L}+\psi}{2\mathbb{L}^2\psi-2\mathbb{L}\psi+1}$ |

Table S11: Consanguinities between different gene positions under haploidy. Under our fixed sib-mating scenario $\psi = s$, under our DMD scenario $\psi = 1 - d_m$, and under the DDM scenario $\psi = (1 - d_f)(1 - d_m)$, $\mathbb{L}$ represents the proportion of paternal transmission.

| | Diploidy | Male PGE | Female MGE |
|---|---|---|---|
| $\rho^{i}_{ff,fm}$ | $\frac{\psi}{4-3\psi}$ | $\frac{\psi}{4-3\psi}$ | $\frac{\psi}{4-3\psi}$ |
| $\rho^{i}_{mf,mm}$ | $\frac{\psi}{4-3\psi}$ | $\frac{\psi}{4-3\psi}$ | $\frac{\psi}{4-3\psi}$ |
| $\rho^{p}_{ff,ff}$ | $\frac{\psi-2}{3\psi-4}$ | $\frac{2-\psi}{4-3\psi}$ | 1 |
| $\rho^{p}_{ff,fm}$ | $\frac{\psi}{4-3\psi}$ | $\frac{\psi}{4-3\psi}$ | $\frac{\psi}{4-3\psi}$ |
| $\rho^{p}_{ff,mf}$ | $\frac{\psi-2}{3\psi-4}$ | $\frac{2-\psi}{4-3\psi}$ | 1 |
| $\rho^{p}_{ff,mm}$ | $\frac{\psi}{4-3\psi}$ | $\frac{\psi}{4-3\psi}$ | $\frac{\psi}{4-3\psi}$ |
| $\rho^{p}_{fm,fm}$ | $\frac{\psi-2}{3\psi-4}$ | 1 | $\frac{2-\psi}{4-3\psi}$ |
| $\rho^{p}_{fm,mf}$ | $\frac{\psi}{4-3\psi}$ | $\frac{\psi}{4-3\psi}$ | $\frac{\psi}{4-3\psi}$ |
| $\rho^{p}_{fm,mm}$ | $\frac{\psi-2}{3\psi-4}$ | 1 | $\frac{2-\psi}{4-3\psi}$ |
| $\rho^{p}_{mf,mf}$ | $\frac{\psi-2}{3\psi-4}$ | $\frac{2-\psi}{4-3\psi}$ | 1 |
| $\rho^{p}_{mf,mm}$ | $\frac{\psi}{4-3\psi}$ | $\frac{\psi}{4-3\psi}$ | $\frac{\psi}{4-3\psi}$ |
| $\rho^{p}_{mm,mm}$ | $\frac{\psi-2}{3\psi-4}$ | 1 | $\frac{2-\psi}{4-3\psi}$ |

Table S12: Consanguinities between different gene positions in offspring under our different genetical systems. Under our fixed sib-mating scenario $\psi = s$, under our labile sib-mating scenario $\psi = 1 - d_m$, and under our viscous population $\psi = (1-d_f)(1-d_m)$. Here we have assumed that in both the male PGE and female MGE scenarios $\mathbb{L} = 0$.

| | Diploidy | Male PGE | Female PGE |
|---|---|---|---|
| $\rho^{Mot}_{ff,ff}$ | $\frac{\psi-2}{3\psi-4}$ | $\frac{\psi-2}{3\psi-4}$ | $-\frac{\psi}{3\psi-4}$ |
| $\rho^{Mot}_{ff,fm}$ | $-\frac{\psi}{3\psi-4}$ | $\frac{(\psi-2)\psi}{3\psi-4}$ | $\frac{(\psi-2)\psi}{3\psi-4}$ |
| $\rho^{Mot}_{ff,mf}$ | $\frac{\psi-2}{3\psi-4}$ | $\frac{\psi-2}{3\psi-4}$ | $-\frac{\psi}{3\psi-4}$ |
| $\rho^{Mot}_{ff,mm}$ | $-\frac{\psi}{3\psi-4}$ | $\frac{(\psi-2)\psi}{3\psi-4}$ | $\frac{(\psi-2)\psi}{3\psi-4}$ |
| $\rho^{Mot}_{fm,ff}$ | $\frac{\psi-2}{3\psi-4}$ | $\frac{\psi-2}{3\psi-4}$ | 1 |
| $\rho^{Mot}_{fm,fm}$ | $-\frac{\psi}{3\psi-4}$ | $-\frac{\psi^2}{3\psi-4}$ | $-\frac{\psi}{3\psi-4}$ |
| $\rho^{Mot}_{fm,mf}$ | $\frac{\psi-2}{3\psi-4}$ | $\frac{\psi-2}{3\psi-4}$ | 1 |
| $\rho^{Mot}_{fm,mm}$ | $-\frac{\psi}{3\psi-4}$ | $-\frac{\psi^2}{3\psi-4}$ | $-\frac{\psi}{3\psi-4}$ |

Table S13: Consanguinities between different gene positions in mothers and offspring under our different genetical systems. Under our fixed sib-mating scenario $\psi = s$, under our labile sib-mating scenario $\psi = 1 - d_m$, and under our viscous population $\psi = (1 - d_f)(1 - d_m)$. Here we have assumed that in both the male PGE and female MGE scenarios $\mathbb{L} = 0$.

| | Diploidy | Male PGE | Female PGE |
|---|---|---|---|
| $\rho$Fomfff | $-\frac{\psi}{3\psi-4}$ | $-\frac{\psi}{3\psi-4}$ | $-\frac{\psi^2}{3\psi-4}$ |
| $\rho$Fomffm | $\frac{\psi-2}{3\psi-4}$ | 1 | $\frac{\psi-2}{3\psi-4}$ |
| $\rho$Fomfmf | $-\frac{\psi}{3\psi-4}$ | $-\frac{\psi}{3\psi-4}$ | $-\frac{\psi^2}{3\psi-4}$ |
| $\rho$Fomfmm | $\frac{\psi-2}{3\psi-4}$ | 1 | $\frac{\psi-2}{3\psi-4}$ |
| $\rho$Fommff | $-\frac{\psi}{3\psi-4}$ | $\frac{(\psi-2)\psi}{3\psi-4}$ | $\frac{(\psi-2)\psi}{3\psi-4}$ |
| $\rho$Fommfm | $\frac{\psi-2}{3\psi-4}$ | $-\frac{\psi}{3\psi-4}$ | $\frac{\psi-2}{3\psi-4}$ |
| $\rho$Fommmf | $-\frac{\psi}{3\psi-4}$ | $\frac{(\psi-2)\psi}{3\psi-4}$ | $\frac{(\psi-2)\psi}{3\psi-4}$ |
| $\rho$Fommmm | $\frac{\psi-2}{3\psi-4}$ | $-\frac{\psi}{3\psi-4}$ | $\frac{\psi-2}{3\psi-4}$ |

Table S14: Consanguinities between different gene positions in fathers and offspring under our different genetical systems. Under our fixed sib-mating scenario $\psi = s$, under our labile sib-mating scenario $\psi = 1 - d_m$, and under our viscous population $\psi = (1 - d_f)(1 - d_m)$. Here we have assumed that in both the male PGE and female MGE scenarios $\mathbb{L} = 0$.

| | $c_{ff}$ | $c_{fm}$ | $c_{mf}$ | $c_{mm}$ | $c_i$ |
|---|---|---|---|---|---|
| Haploidy | $\frac{1}{4}$ | $\frac{1}{4}$ | $\frac{1}{4}$ | $\frac{1}{4}$ | N/A |
| Diploidy | $\frac{1}{3}$ | $\frac{1}{3}$ | $\frac{1}{3}$ | N/A | N/A |
| Arrhenotoky | N/A | $\frac{1}{3}$ | $\frac{1}{3}$ | $\frac{1}{3}$ | N/A |
| Paterothylotoky | $\frac{\mathbb{L}-1}{2\mathbb{L}-3}$ | $\frac{\mathbb{L}-1}{2\mathbb{L}-3}$ | $\frac{\mathbb{L}-1}{2\mathbb{L}-3}$ | $\frac{\mathbb{L}}{3-2\mathbb{L}}$ | N/A |
| Male PGE | $\frac{\mathbb{L}}{3-2\mathbb{L}}$ | $\frac{\mathbb{L}-1}{2\mathbb{L}-3}$ | $\frac{\mathbb{L}-1}{2\mathbb{L}-3}$ | $\frac{\mathbb{L}-1}{2\mathbb{L}-3}$ | N/A |
| Female PGE | $\frac{1}{4}$ | $\frac{1}{4}$ | $\frac{1}{4}$ | $\frac{1}{4}$ | N/A |
| Icerya system | $\frac{\varphi-1}{2\varphi-3}$ | $\frac{\varphi-1}{2\varphi-3}$ | $\frac{\varphi-1}{2\varphi-3}$ | 0 | $\frac{\varphi}{3-2\varphi}$ |

Table S15: Class reproductive values for the gene positions in various genetic systems. Where $c_{ff}$ are female maternal-origin genes, $c_{fm}$ are female paternal-origin genes, $c_{mf}$ are male maternal-origin genes, $c_{mm}$ are male paternal-origin genes, and $c_i$ is the infectious male lineage

positions (Taylor, 1990; Taylor, 1996). Solving for our different genetic systems we get the class reproductive values seen in Table S15.

$$\begin{pmatrix} c_{ff} & c_{fm} & c_{mf} & c_{mm} \end{pmatrix} = \begin{pmatrix} c_{ff} & c_{fm} & c_{mf} & c_{mm} \end{pmatrix} \begin{pmatrix} \pi_{ff,ff} & \pi_{ff,fm} & \pi_{ff,mf} & \pi_{ff,mm} \\ \pi_{fm,ff} & \pi_{fm,fm} & \pi_{fm,mf} & \pi_{fm,mm} \\ \pi_{mf,ff} & \pi_{mf,fm} & \pi_{mf,mf} & \pi_{mf,mm} \\ \pi_{mm,ff} & \pi_{mm,fm} & \pi_{mm,mf} & \pi_{mm,mm} \end{pmatrix} \tag{S3.6.1}$$

##### 3.7 Condition for increase

Putting the reproductive values, consanguinities, and marginal fitness effects together, we can write the condition for increase in our diploid systems with offspring control as so:

$$\frac{\tau}{\sigma} < \frac{\alpha}{\beta}\left(\frac{\mathscr{X} w_{ff} + (1-\mathscr{X}) w_{fm}}{(1-\mathscr{Y}) w_{mf} + \mathscr{Y} w_{mm}}\right) \tag{S3.7.1}$$

Where $w_{ff}$ is the inclusive fitness (IF) effect to a female maternal-origin copy of an increase in the trait value, $w_{fm}$ is the IF effect to a female paternal-origin gene copy of a change in the trait value, $w_{mf}$ is the IF effect to a male maternal-origin gene copy of a change in the trait value, and $w_{mm}$ is the IF effect to a male paternal-origin gene copy of a change in the trait value. The $\alpha$,$\beta$,$\mathscr{X}$, and $\mathscr{Y}$, scale the relative phenotypic effects of these gene copies on males and females.

Where the inclusive fitness effects experienced by our different genetic actors is:

$$\begin{aligned} w_{ff} =& c_{fm}\left(\rho^{i}_{ff,fm} - a_f \rho^{p}_{ff,fm}\right) + c_{ff}\left(1 - a_f \rho^{p}_{ff,ff}\right) \\ &+ c_{mm}\left(\psi - a_{mf}\right)\rho^{p}_{ff,mm} + c_{mf}\left(\psi - a_{mf}\right)\rho^{p}_{ff,mf} \end{aligned} \tag{S3.7.2a}$$

$$\begin{aligned} w_{fm} =& c_{ff}\left(\rho^{i}_{ff,fm} - a_f \rho^{p}_{ff,fm}\right) + c_{fm}\left(1 - a_f \rho^{p}_{fm,fm}\right) \\ &+ c_{mm}\left(\psi - a_{mf}\right)\rho^{p}_{fm,mm} + c_{mf}\left(\psi - a_{mf}\right)\rho^{p}_{fm,mf} \end{aligned} \tag{S3.7.2b}$$

$$w_{mf} = c_{mm}\left(\rho^{i}_{mf,mm} - a_{mm}\rho^{p}_{mf,mm}\right) + c_{mf}\left(1 - a_{mm}\rho^{p}_{mf,mf}\right) \tag{S3.7.2c}$$

$$w_{mm} = c_{mf}\left(\rho^{i}_{mf,mm} - a_{mm}\rho^{p}_{mf,mm}\right) + c_{mm}\left(1 - a_{mm}\rho^{p}_{mm,mm}\right) \tag{S3.7.2d}$$

#### 3.8 Trait Optima

To find the optimal trait level in our toy model, we set our condition for increase to zero, and then solve for $\phi^*$. In the main text this is referred to as $z^*$.

| | Trait optima $\phi^*$ |
|---|---|
| Haploidy | $\frac{\hat{z}_f \mathrm{SD}_m^2\left((2(\mathbb{L}-1)\mathbb{L}+1)\left(d_f-2\right)d_f(\mathbb{L}(s-1)+1)+2\mathbb{L}(\mathbb{L}-1)^2(s-1)\right)+\mathbb{L}(s-1)\mathrm{SD}_f^2\hat{z}_m}{(2(\mathbb{L}-1)\mathbb{L}+1)\left(d_f-2\right)d_f\mathrm{SD}_m^2(\mathbb{L}(s-1)+1)+\mathbb{L}(s-1)\left(\mathrm{SD}_f^2+2(\mathbb{L}-1)^2\mathrm{SD}_m^2\right)}$ |
| Arrhenotoky/MPGE | $\frac{\alpha\hat{z}_f\mathrm{SD}_m^2\left(\left(d_f-2\right)d_f((s-2)s(2\mathscr{X}-1)+2(\mathscr{X}-2))-2(s-1)\mathscr{X}\right)-\beta(s-1)\mathrm{SD}_f^2\hat{z}_m(2(s-2)\mathscr{Y}-s+4)}{\alpha^2\mathrm{SD}_m^2\left(\left(d_f-2\right)d_f((s-2)s(2\mathscr{X}-1)+2(\mathscr{X}-2))-2(s-1)\mathscr{X}\right)-\beta^2(s-1)\mathrm{SD}_f^2(2(s-2)\mathscr{Y}-s+4)}$ |
| Paterothylotoky/FMGE | $\frac{\alpha\hat{z}_f\mathrm{SD}_m^2\left(\left(d_f-2\right)d_f\left(2(s-1)^2\mathscr{X}-s-2\right)+2(s-1)(\mathscr{X}-1)\right)-2\beta(s-1)\mathrm{SD}_f^2\hat{z}_m(s(\mathscr{Y}-1)+2)}{\alpha^2\mathrm{SD}_m^2\left(\left(d_f-2\right)d_f\left(2(s-1)^2\mathscr{X}-s-2\right)+2(s-1)(\mathscr{X}-1)\right)-2\beta^2(s-1)\mathrm{SD}_f^2(s(\mathscr{Y}-1)+2)}$ |
| Diploidy | $\frac{\alpha\hat{z}_f\mathrm{SD}_m^2\left((s+1)\left(d_f-2\right)d_f+s-1\right)+2\beta(s-1)\mathrm{SD}_f^2\hat{z}_m}{\alpha^2\mathrm{SD}_m^2\left((s+1)\left(d_f-2\right)d_f+s-1\right)+2\beta^2(s-1)\mathrm{SD}_f^2}$ |

Table S16: Optima for a normally distributed trait when under offspring control, and there is sib-mating

| | Trait optima $\phi^*$ |
|---|---|
| Haploidy | $\frac{\left(d_f-2\right)d_f\hat{z}_f\mathrm{SD}_m^2(\mathbb{L}(s-1)+1)+\mathbb{L}(s-1)\mathrm{SD}_f^2\hat{z}_m}{\left(d_f-2\right)d_f\mathrm{SD}_m^2(\mathbb{L}(s-1)+1)+\mathbb{L}(s-1)\mathrm{SD}_f^2}$ |
| Arrhenotoky/MPGE | $\frac{\left(d_f-2\right)d_f\hat{z}_f\mathrm{SD}_m^2(2(s-1)s\mathscr{X}-s-2)+(s-2)(s-1)\mathrm{SD}_f^2\hat{z}_m}{\alpha\left(d_f-2\right)d_f\mathrm{SD}_m^2(2(s-1)s\mathscr{X}-s-2)+\alpha(s-2)(s-1)\mathrm{SD}_f^2}$ |
| Paterothylotoky/FMGE | $\frac{s(2s-5)\left(d_f-2\right)d_f\hat{z}_f\mathrm{SD}_m^2+2(s-2)(s-1)\mathrm{SD}_f^2\hat{z}_m}{\alpha s(2s-5)\left(d_f-2\right)d_f\mathrm{SD}_m^2+2\alpha(s-2)(s-1)\mathrm{SD}_f^2}$ |
| Diploidy | $\frac{(s+1)\left(d_f-2\right)d_f\hat{z}_f\mathrm{SD}_m^2+(s-1)\mathrm{SD}_f^2\hat{z}_m}{\alpha(s+1)\left(d_f-2\right)d_f\mathrm{SD}_m^2+\alpha(s-1)\mathrm{SD}_f^2}$ |

Table S17: Optima for a normally distributed trait when under maternal control, and there is sib-mating

| | Trait optima $\phi^*$ |
|---|---|
| Haploidy | $\frac{\left(d_f-2\right)d_f\hat{z}_f\mathrm{SD}_m^2(\mathbb{L}(s-1)+1)+\mathbb{L}(s-1)\mathrm{SD}_f^2\hat{z}_m}{\left(d_f-2\right)d_f\mathrm{SD}_m^2(\mathbb{L}(s-1)+1)+\mathbb{L}(s-1)\mathrm{SD}_f^2}$ |
| Arrhenotoky/MPGE | $\frac{((s-2)s+4)\left(d_f-2\right)d_f\hat{z}_f\mathrm{SD}_m^2+(s-1)s\mathrm{SD}_f^2\hat{z}_m}{\beta((s-2)s+4)\left(d_f-2\right)d_f\mathrm{SD}_m^2+\beta(s-1)s\mathrm{SD}_f^2}$ |
| Paterothylotoky/FMGE | $\frac{\left(d_f-2\right)d_f\hat{z}_f\mathrm{SD}_m^2(s((s-1)s(2\mathscr{Y}-1)-1)-2)+(s-1)\mathrm{SD}_f^2\hat{z}_m((s-1)s(2\mathscr{Y}-1)-2)}{\beta\left(\left(d_f-2\right)d_f\mathrm{SD}_m^2(s((s-1)s(2\mathscr{Y}-1)-1)-2)+(s-1)\mathrm{SD}_f^2((s-1)s(2\mathscr{Y}-1)-2)\right)}$ |
| Diploidy | $\frac{(s+1)\left(d_f-2\right)d_f\hat{z}_f\mathrm{SD}_m^2+(s-1)\mathrm{SD}_f^2\hat{z}_m}{\beta(s+1)\left(d_f-2\right)d_f\mathrm{SD}_m^2+\beta(s-1)\mathrm{SD}_f^2}$ |

Table S18: Optima for a normally distributed trait when under paternal control, and there is sib-mating
